# SREBP2 delivery to striatal astrocytes normalizes transcription of cholesterol biosynthesis genes and ameliorates pathological features in Huntington’s Disease

**DOI:** 10.1101/2020.11.23.393793

**Authors:** Giulia Birolini, Gianluca Verlengia, Francesca Talpo, Claudia Maniezzi, Lorena Zentilin, Mauro Giacca, Paola Conforti, Chiara Cordiglieri, Claudio Caccia, Valerio Leoni, Franco Taroni, Gerardo Biella, Michele Simonato, Elena Cattaneo, Marta Valenza

## Abstract

Brain cholesterol is produced mainly by astrocytes and is important for neuronal function. Its biosynthesis is severely reduced in mouse models of Huntington’s Disease (HD). One possible mechanism is a diminished nuclear translocation of the transcription factor sterol regulatory element binding protein 2 (SREBP2) and, consequently, reduced activation of SREBP-controlled genes in the cholesterol biosynthesis pathway.

Here we evaluated the efficacy of a gene therapy based on the unilateral intra-striatal injection of a recombinant adeno-associated virus 2/5 (AAV2/5) targeting astrocytes specifically and carrying the N-terminal fragment of human SREBP2 (hSREBP2).

Robust hSREBP2 expression in striatal glial cells in HD mice activated the transcription of cholesterol biosynthesis pathway genes, restored synaptic transmission, reversed *Drd2* transcript levels decline, cleared muHTT aggregates and attenuated behavioral deficits. We conclude that glial SREBP2 participates in HD brain pathogenesis in vivo and that AAV-based delivery of SREBP2 to astrocytes counteracts key features of HD.

## Introduction

Cholesterol is a multifaceted molecule that plays key roles in the brain during development and in adulthood. Its concentration in the brain is higher than in any other tissue (15–20 mg/g tissue)^1^. Up to 70–80% of cholesterol in the adult brain is present in myelin sheaths, while the rest is localized in the plasma membranes of astrocytes and neurons^2^. In neurons cholesterol plays important roles in synaptic transmission, as it is required for synaptic vesicle formation and function^3,4,5^ and for optimal neurotransmitter release^6,7,8^. As the blood-brain barrier (BBB) efficiently prevents the passage of circulating cholesterol, brain cholesterol is synthesized locally^9^. Of note, in adulthood astrocytes are believed to synthetize most cholesterol, which is transferred to the neurons via ApoE-containing lipoproteins^3,10^.

Cholesterol biosynthesis is regulated by sterol regulatory element binding protein 2 (SREBP2), the master transcription factor that activates the expression of most cholesterol biosynthesis genes. When the cholesterol level is low, SREBP2 is cleaved by SREBP cleavage activation protein (SCAP), leading to SREBP2 activation and translocation from the endoplasmic reticulum (ER) to the Golgi apparatus. Within the Golgi apparatus, SREBP2 is cleaved by two proteases called the site 1 (S1P) and site 2 (S2P) proteases, resulting in an active 68-kDa N-terminal fragment that moves to the nucleus, binds the sterol response element (SRE) in the promoters of target genes, and activates their transcription^11^. When cells do not need to produce cholesterol, SREBP2 is retained in the ER by the insulin-induced gene 1 (INSIG1) and insulin-induced gene 2 (INSIG2) proteins^12^.

Dysregulation of brain cholesterol homeostasis has been linked to several neurodegenerative diseases^13,14^. Among these conditions is Huntington’s disease (HD), an adult-onset disorder characterized by motor, cognitive, and psychiatric features. HD is caused by a CAG expansion in the HTT(IT15) gene encoding the huntingtin (HTT) protein and it is characterized by progressive loss of striatal medium-sized spiny neurons (MSNs) and cortical pyramidal neurons^15,16^.

A large number of studies conducted in different HD rodent models have demonstrated significantly reduced biosynthesis of cholesterol in the brain, with the striatum being affected first^17,18,19,20,21^. In the adult HD mouse brain, this dysfunction is evidenced by a reduced rate of de novo cholesterol biosynthesis, precedes the onset of motor symptoms, is CAG-dependent, as confirmed by studies in an allelic series of HD mice, and leads to a reduced content of total cholesterol (and of some of its metabolites and catabolites) at late symptomatic stages^21^. The underlying molecular mechanism implicates reduced activity of SREBP2 in the presence of mutant HTT^17,22^, likely through sequestration in the cytoplasm of the SREBP2/importin β complex required for nuclear import^23^.

Astrocytes are central in this dysfunction, as highlighted by in vitro studies. In fact, SREBP2 gene silencing in wild-type (wt) astrocytes has detrimental effects on HD neurons^22^, while the forced expression of the N-terminal active fragment of SREBP2 in HD astrocytes reverses neurite outgrowth and synaptic defects in HD neurons^22^. Despite these findings, the impact of reduced astrocyte-mediated cholesterol synthesis in vivo in the HD brain and the therapeutic potential of its reversal are not yet documented.

To gain mechanistic insights and confidence in the therapeutic potential of targeting this pathway *in vivo*, we forced the expression of the N-terminal active fragment of human SREBP2 (hSREBP2) in striatal astrocytes of a transgenic HD mouse model by using a recombinant adeno-associated virus 2/5 (AAV2/5). We found that increasing hSREBP2 level specifically in the nucleus of astrocytes stimulates cholesterol biosynthesis in the striatum of HD mice. As a consequence, synaptic transmission of both inhibitory and excitatory synapses is restored, the number of striatal MSNs expressing Drd2 receptors normalized, mutant HTT (muHTT) aggregation reduced, motor defects ameliorated, and cognitive decline is completely rescued.

## Materials and Methods

### Colony management

Our R6/2 colony lifespan was approximately of 13 weeks and it was maintained through the male line exclusively^26^. Transgenic R6/2 males were paired with non-carrier females (B6CBAF1/J, purchased from Charles River). The CAG repeat length of the animals used in this study is 130-170 CAGs (Laragen). Changes that could affect strain productivity, general behavior, litter size, pup survival, genotype frequency, and phenotype were constantly monitored.

The Drd2-eGFP colony is a transgenic mouse line generated in 2003 by the GENSAT (Gene Expression Nervous System Atlas) project at Rockefeller University in New York^60^. Primary labelling was found to occur in Drd2 positive medium spiny neurons from the indirect basal ganglia pathway. In this work, this model was crossed with R6/2 mice in order to label *in vivo* Drd2-expressing medium spiny neurons.

All mice were weaned at 21 day (+/− 3 days). Mice were housed under standard conditions (22 ± 1°C, 60% relative humidity, 12 hours light/dark cycle, 3–4 mice/cage, with food and water provided ad libitum). After PCR genotyping^26,60^, male and female mice were included and randomly divided into experimental groups. Littermates were included as controls.

Animal care was conducted in accordance with standard ethical guidelines approved by the Italian Governing Law (D.lgs 26/2014; Authorization n.324/2015-PR issued May 6, 2015 by Ministry of Health); the NIH Guide for the Care and Use of Laboratory Animals (2011 edition) and the EU directives and guidelines (EEC Council Directive 2010/63/UE) and the local ethics committee approved the experiments.

### Vector production

We obtained the AAV vector plasmid pZac-gfaABC1D-hSREBP2-tdTomato, expressing the active N-terminal fragment of hSREBP2 (1-482 aa) fused with tdTomato, starting from pZac2.1 gfaABC1D-tdTomato (from Addgene) and pcDNA3.1 hSREBP2(402)-eGFP (from T. Osborne)^61^. In the AAV vector plasmids, the transgenes are under the control of the minimal GFAP promoter (gfaABC1D) and are surrounded by the inverted terminal repeats derived from AAV serotype 2.

These plasmids were used to generate the corresponding AAV vectors in the AAV Vector Unit at the International Centre for Genetic Engineering and Biotechnology Trieste (http://www.icgeb.org/avu-core-facility.html), as described previously^62^ with few modifications. Briefly, infectious recombinant AAV vector particles were generated in HEK293T cells culture in roller bottles by a cross-packaging approach whereby the vector genome was packaged into AAV capsid serotype-5. Viral stocks were obtained by PEG precipitation and CsCl2 gradient centrifugation^63^. The physical titer of recombinant AAVs was determined by quantifying vector genomes (vg) packaged into viral particles, by real-time PCR against a standard curve of a plasmid containing the vector genome^64^. Values obtained were 1,7 × 10^14^ U/mL for AAV2/5-gfaABC1D-tdTomato and 1,7 × 10^13^ U/mL for AAV2/5-gfaABC1D-hSREBP2-tdTomato.

HSV1/JDNI8-gfaABC1D-tdTomato (5 × 10^9^ pfu/mL) was produced and purified as previously described^65^. Briefly, ICP4/ICP27-complementing U2OS cells were transfected with a purified BAC-DNA engineered to contain the HSV1-JDNI8/gfaABC1D-tdTomato vector genome. 10 days upon transfection, the virus-containing supernatant was collected and used for subsequent infections of ICP4/ICP27-U2OS complementing cells in order to obtain a highly-concentrated monoclonal vector stock suitable for in vivo applications. Estimation of HSV1 vector titers was carried out by qPCR quantification of glycoprotein D gene copy number^65^.

### Stereotaxic injection of HSV1/JDNI8-gfaABC1D-tdTomato, AAV2/5-gfaABC1D-tdTomato and AAV2/5-gfaABC1D-hSREBP2-tdTomato

7-weeks-old mice were deeply anesthetized with Avertin 2.5% (15 *μ*L/gr body weight). The virus was injected by implantation of a borosilicate glass needle into the right striatum of mice via stereotaxic surgery using an automated infusion syringe pump (KD Scientific, KDS100), on which a 50 *μ*L Gastight Syringe Model 1705 TLL with PTFE Luer Lock (Hamilton, 80920) could be mounted. In order to favor needle entry and vector spread in the injected striata, we used borosilicate needles customized by laser shaping with the Leica Laser Microdissector CTR6000 (Leica Microsystems), that allowed a 45 degrees edge chamfering of tip (inner diameter at tip = 60 *μ*m) and, moreover, to open an additional circular hole (Ø 20 *μ*m) about 100 *μ*m from the beveled edge^66^.

The following stereotaxic coordinates were used: 2 mm lateral to midline, 0.74 mm rostral to the bregma, 3.5 mm ventral to the skull surface; from Paxinos G and Watson C. The Rat Brain in Stereotaxic Coordinates. Academic Press, San Diego. The rate of injection was 12 *μ*l/min with a total volume of 2 *μ*L.

Assessment of post-operative pain and distress was performed using a specific table for pain scoring based on behavioral indicators of well-being and monitoring mice body weight^67^.

### Immunohistochemistry analysis

Four weeks after the infection anesthetized mice were transcardially perfused with PFA 4%. Brains were post-fixed overnight in PFA 4% at 4°C and then in 30% sucrose to prevent ice crystal damage during freezing in OCT.

15 μm coronal sections or 30 μm coronal sections were prepared for immunohistochemical analysis. Epitopes were demasked at 98°C with NaCitrate 10 mM and then slices were incubated with the following primary antibodies for 3h at RT: rabbit anti-SREBP2 (1:100; Ls-Bio, LS-C179708), rabbit anti-RFP (1:100, MBL, PM005), mouse anti-RFP (1:100, Thermo Fisher, MA5-15257), anti-DARPP32 (1:100; Cell Signalling, 2306), mouse anti-NeuN (1:100; Millipore, MAB377); rabbit anti-GFAP (1:250; Dako, Z0334), rabbit anti-IBA1 (1:100, Wako, 019-19741), rabbit anti-Huntingtin clone EM48 (1:100; Millipore, MAB5374). Anti-rabbit Alexa Fluor 568-conjugated goat secondary antibodies (1:500; Invitrogen), anti-rabbit Alexa Fluor 633-conjugated goat secondary antibodies (1:500; Invitrogen) or anti-mouse Alexa Fluor 488-conjugated goat secondary antibodies (1:500; Invitrogen) were used for detection (1h at RT). Sections were counterstained with the nuclear dye Hoechst 33258 (1:10.000, Invitrogen) and then mounted under cover slips using Vectashield (Vector Laboratories).

### Image acquisition and quantification

To study the spread of the viruses, large images were acquired at 4× magnification (NA 0.75) with a customized Nikon Ti microscope, equipped with a X-light V2 spinning-disk scan head with VCS module for structure illumination (Crest Optics).

To study the tropism of the viruses and to count muHTT aggregates, confocal images (5 to 10-z steps) were acquired with a LEICA SP5 laser scanning confocal microscope. Laser intensity and detector gain were maintained constant for all images. To count aggregates in the striatum 18 images/mice from 9 sections throughout the entire striatum were taken from three R6/2-hBP2 mice at 40× magnification (NA 1.40). To quantify the number and the size of aggregates, ImageJ software (on the Fiji platform) was used to measure fluorescence signal. Images were split into three-color channels and the same global threshold was set on each signal histogram.

### RNA extraction and qRT-PCR

Four weeks after the infection, mice were sacrificed by cervical dislocation and tissues were isolated and frozen. Total RNA from the infused striatum and the ipsi-lateral cortex of treated mice was extracted with TRIzol reagent (Life Technologies, 15596026). RNA quality check was carried out on 1% agarose gel. Potential contamination of DNA was removed using the Ambion^®^ DNA-free™ Kit (Invitrogen, AM1906). 500 ng of RNA was retrotranscribed to single-stranded cDNA using the iScript cDNA synthesis kit (Bio-Rad, 1708891). The analyses were performed in 3 animals/group. For each sample, two different reverse transcription were performed. For each cDNA reaction, one or two conventional qPCR were performed in a CFX96 Real-Time System (Bio-Rad) in a 15 *μ*l volume containing diluited cDNA and 7.5 μL iQ EVA Green Supermix (Bio-Rad, 172-5204), 7.5 μL nuclease-free water and 0.33 μM forward and reverse primers. The amounts of target transcript were normalized to β-actin as reference gene. Table S1 summarizes the primer sequences and melting temperature used in this work.

### Mass Spectrometry analysis

To a screw-capped vial sealed with a Teflon-lined septum were added 50 μL of homogenates together with 500 ng of D4-lathosterol (CDN Isotopes, Canada), 500 ng of D6-desmosterol (Avantipolar Lipids, USA), 100 ng of D6-lanosterol (Avantipolar Lipids, USA), 400 ng of D7-24S-hydroxycholesterol (Avantipolar Lipids, USA), and 100 μg of epicoprostanol (Sigma-Merck) as internal standards, 50 *μ*L of butylated hydroxytoluene (BHT) (5 g/L) and 25 *μ*L of EDTA (10 g/L). Argon was flushed through to remove air. Alkaline hydrolysis was allowed to proceed at room temperature (22°C) for 1h in the presence of 1 M ethanolic potassium hydroxide solution under magnetic stirring. After hydrolysis, the neutral sterols (cholesterol, lathosterol, desmosterol and lanosterol) and 24S-OHC were extracted twice with 5ml of hexane. The organic solvents were evaporated under a gentle stream of nitrogen and converted into trimethylsilyl ethers with BSTFA-1% TMCS (Cerilliant, USA) at 70 °C for 60 min. Analysis was performed by gas chromatography - mass spectrometry (GC–MS) on a Clarus 600 gas chromatograph (Perkin Elmer, USA) equipped with Elite-5MS capillary column (30 m, 0.32 mm, 0.25μm. Perkin Elmer, USA) connected to Clarus 600C mass spectrometer (Perkin Elmer, USA). The oven temperature program was as follows: initial temperature 180 °C was held for 1 min, followed by a linear ramp of 20 °C/min to 270 °C, and then a linear ramp of 5 °C/min to 290 °C, which was held for 10 min. Helium was used as carrier gas at a flow rate of 1 mL/min and 1μL of sample was injected in splitless mode. Mass spectrometric data were acquired in selected ion monitoring mode. Peak integration was performed manually. Sterols and 24S-OHC were quantified against internal standards, using standard curves for the listed sterols^68^.

### Synaptosomes preparation and WB analysis

Mice were sacrificed by cervical dislocation and the tissues were isolated and frozen. Syn-PER Synaptic Protein Extraction Reagent (Thermo Fisher Scientific, 87793) was used for synaptosome purification from the striatum and the ipsi-lateral cortex accordingly to the manufacturer instruction. Briefly, a volume of 10 mL of Syn-PER Reagent per gram of tissue was added and tissues were homogenized on ice. The homogenate was centrifuged at 1,200 x g for 10 min at 4°C. The supernatant was centrifuged at 15,000 x g for 20 min at 4°C to pellet synaptosomes. The pellet was resuspended in Syn-PER Reagent. Proteins were quantified with Pierce™ BCA Protein Assay Kit (Thermo Fisher Scientific, 23225).

Criterion TGX Stain Free Precast Gels 7.5% (Bio-Rad, 5678023) or Any kD (Bio-Rad, 5678123) were used for Western Blot analysis. Membranes were incubated with the following primary antibodies over night at 4°C: rabbit anti-RFP (1:1000; MBL, PM005), rabbit anti-SREBP2 (1:1000; LS-Bio, LS-B4695), mouse anti-SYP (1:500; Abcam, ab8049), mouse anti-SNAP25 (1:1000; Abcam, ab66066), rabbit anti-VAMP1 (1:1000; Abcam, ab151712), mouse anti-PSD95 (1:1000; SySy, 124011), rabbit anti-SHANK3 (1:1000; SySy, 162302), rabbit anti-NMDAR1 (1:500; MerckMillipore, Ab9864), rabbit anti-GAPDH (1:5000; Abcam, ab37168). Goat anti-mouse IgG-HRP (1:3000; Bio-Rad, 1706516) or Goat anti-rabbit IgG-HRP (1:3000; Bio-Rad, 1706515) were used for detection (1h at RT). Proteins were detected using Clarity Western ECL Substrate (Bio-Rad, 1705061) or SuperSignal West Femto Maximum Sensitivity Substrate (Thermo Fisher Scientific, 34096) using ChemiDoc MP System (Bio-Rad). Protein levels were normalized using GAPDH (for the cytosolic fraction) or using 6 different protein bands of the stain-free technology (for synaptosomes). 4 animals/group were analyzed by western blotting and for each protein, two or three technical replicates were performed.

### X-CLARITY processing and 2-photon imaging of cleared tissues

4 weeks after the infection anesthetized mice were transcardially perfused with PFA 4%. Brains were post-fixed overnight and sliced with a Leica VT1000S Vibrating blade microtome (Leica Biosystems). From each mouse, two 1-mm thick coronal brain slices were prepared, from which the infused and the contra-lateral striatum were isolated. Tissues were clarified using the X-CLARITY™ system (Logos Biosystem) according to manufacturer instructions. Briefly, tissues were incubated in embedding solution (Logos Biosystems, C13104) at 4°C for 24h at most, allowing hydrogel monomers to diffuse uniformly throughout the samples and to covalently link biomolecules including proteins, nucleic acids and small molecules. Polymerization was performed by placing the samples within the X-CLARITY™ Polymerization at 37°C for 3 hours under vacuum condition (#x2212;90kPa). Following washing steps, the hydrogel-embedded tissues were rinsed with Electrophoretic Tissue Clearing (ETC) Solution (Logos Biosystems, C13001) and moved into the X-CLARITY™ Tissue Clearing System II (Logos Biosystems, C30001) chamber, filled with the ETC solution in which the application of a uniform electric current (1.5 A) at a controlled temperature of 37°C enabled the active extraction of lipids from tissues in about 2-3 hours.

After clearing, samples were washed with PBS 1X overnight at RT to remove residual SDS. The endogenous signal (eGFP) was acquired by using a A1 MP+ microscope (Nikon) equipped with Ti:Sapphire MP tunable laser (680 - 1080 nm) using the water plan-Apo LWD 25 X objective (NA 1.1). Images were acquired with the following settings: 16 bit, Z-stack of approximately 400 μm with a Z step size of 5 μm. The endogenous fluorescent protein eGFP was excited at a wavelength of 900 nm. For each striatum, 2-3 images were acquired. Images were processed using NIS-Elements AR software v5.21(Nikon-Lim). First, a 3D deconvolution algorithm (Richardson-Lucy with 20 itations) was applied to improve image quality. Afterwards, background was removed using the rolling ball correction. To quantify the number of cells expressing eGFP, a segmentation 3D-object mask was applied over the 3D-reconstructed volume, using a specialized algorithm package in NIS-Elements AR v5.21. For each image, the number of cells expressing eGFP was normalized on the Z volume acquired.

### Electrophysiological analysis

Experiments were performed on submerged brain slices obtained from 12-week-old mice. Animals were anesthetized by inhalation of isoflurane and decapitated. The head was rapidly submerged in ice-cold (~ 4°C) and oxygenated (95% O_2_ - 5% CO_2_) cutting solution containing: Sucrose 70 mM, NaCl 80 mM, KCl 2.5 mM, NaHCO_3_ 26 mM, Glucose 15 mM, MgCl_2_ 7 mM, CaCl_2_ 1 mM and NaH_2_PO_4_ 1.25 mM. Striatal coronal slices (300-μm-thick) were cut using a vibratome (DTK-1000, Dosaka EM, Kyoto, Japan) and allowed to equilibrate for at least 1 hour in a chamber filled with oxygenated ACSF containing: NaCl 125 mM, KCl 2.5 mM, NaHCO_3_ 26 mM, Glucose 15 mM, MgCl_2_ 1.3 mM, CaCl_2_ 2.3 mM and NaH_2_PO_4_ 1.25 mM. The slices collected from the hemisphere ipsilateral to the infusion site were transferred to a submerged-style recording chamber at room temperature (~ 23-25°C) and were continuously perfused at 1.4 ml/min with ACSF. The chamber was mounted on an E600FN microscope (Nikon) equipped with 4X and 40X water-immersion objectives (Nikon) and connected to a near-infrared CCD camera for cells visualization.

Data were obtained from MSNs using the whole-cell patch-clamp technique in both voltage-and current-clamp mode. The patch pipette was produced from borosilicate glass capillary tubes (Hilgenberg GmbH) using a horizontal puller (P-97, Sutter instruments) and filled with an intracellular solution containing: Cs-methanesulphonate 120 mM, KCl 5 mM, CaCl_2_ 1 mM, MgCl_2_ 2 mM, EGTA 10 mM, Na_2_ATP 4 mM, Na_3_GTP 0.3 mM, Hepes 8 mM and lidocaine N-ethylbromide 5 mM (added to inhibit firing by blocking intracellularly the voltage-sensitive Na^+^ channels) (pH adjusted to 7.3 with KOH). Spontaneous excitatory postsynaptic currents (sEPSCs), mediated by the activation of ionotropic glutamate receptors, were recorded from MSNs at a holding potential of −70 mV, whereas spontaneous inhibitory postsynaptic currents (sIPSCs), elicited by the activation of GABA_A_ receptors, were assessed at a holding potential of 0 mV. The signals were amplified with a MultiClamp 700B amplifier (Molecular Devices) and digitized with a Digidata 1322 computer interface (Digitata, Axon Instruments Molecular Devices, Sunnyvale, CA). Data were acquired using the software Clampex 9.2 (Molecular Devices, Palo Alto, CA, U.S.A.), sampled at 20 kHz and filtered at 2 kHz.

The off-line detection of spontaneous postsynaptic currents (sPSCs) were performed manually using a custom-made software in Labview (National Instruments, Austin, TX, U.S.A.). The amplitudes of sPSCs obeyed a lognormal distribution. Accordingly, the mean amplitude was computed as the peak of the lognormal function used to fit the distribution. Intervals (measured as time between two consecutive sPSCs) for spontaneous events were distributed exponentially and the mean interval was computed as the tau (τ_interval_) value of the mono-exponential function that best fitted this distribution. The reciprocal of τ (1/τ) is the mean of the instantaneous frequencies of sPSCs. Furthermore, the analysis of the membrane capacitance (C_m_) and the input resistance (R_in_) was performed using Clampfit 10.2 (Molecular Devices, Palo Alto, CA, U.S.A.). C_m_ was estimated from the capacitive current evoked by a −10 mV pulse, whereas R_in_ was calculated from the linear portion of the I-V relationship obtained by measuring steady-state voltage responses to hyperpolarizing and depolarizing current steps.

### Behavioral tests

Mice behavior was evaluated at 11 weeks of age. Animals were assigned randomly and sex was balanced in the various experimental groups. All the behavioral analyses were performed in blind.

#### Activity Cage

spontaneous locomotor activity was evaluated by the activity cage test, in presence of a low-intensity white light source. The animal was placed in the center of the testing, transparent, arena (25 cm × 25 cm) (2Biological Instrument) and allowed to freely move for an hour. Following 15 minutes of habituation, both horizontal and vertical motor activities were assessed by an automated tracking system (Actitrack software, 2Biological Instrument) connected to infrared sensors surrounding the arena. Total distance travelled, mean velocity speed, stereotyped movements and numbers of rearings were evaluated. The % of time that mice explored the periphery or the center area of the was evaluated as a measure of anxiety-like behavior.

#### Novel Object Recognition (NOR) test

long-term memory was evaluated by the NOR test, using a grey-colored, non-reflective arena (44 × 44 × 44 cm). All phases of the test were conducted with a low-intensity white light source. In a first habituation phase, mice were placed into the empty arena for 10 min. The habituation phase was followed by the familiarization one, in which two identical objects (A′ and A″) were presented to each animal for 10 min. Twenty-four hours later, during the testing phase, the same animals were exposed to one familiar object (A″) and a new object (B) for 10 min. A measure of the spontaneous recognition memory was represented by the index of discrimination, calculated as (time exploring the novel object − time exploring the familiar object) / (time exploring both objects) × 100. Mice exploring less than 7 sec. were excluded from the analysis due to their inability to perform the task.

#### Grip strength test

mice were lifted by the lower back and tail and lowered towards the grip (Ugo Basile) until the animal grabbed it with both front paws. The animal was then lowered toward the platform and gently pulled straight back with consistent force until it released its grip. The forelimb grip force, measured in grams, was recorded. The test was repeated for 5 times, and measures were averaged. After testing, animals were placed back into their home cage.

#### Paw clasping test

mice were suspended by the tail for 30 s and the clasping phenotype was graded according to the following scale: level 0, no clasping; level 1, one hindlimb retracted toward the abdomen; level 2, both hindlimbs retracted toward the abdomen; level 3, both hindlimbs entirely retracted and touching the abdomen. After testing, animals were placed back into their home cage.

### Statistics

Prism 8 (GraphPad software) was used to perform all statistical analyses. Data are presented as means ± standard error of the mean (SEM). Grubbs’ test was applied to identify outliers. For each set of data to be compared, we determined whether data were normally distributed or not to select parametric or not parametric statistical tests. The specific statistical test used is indicated in the legend of all results figures. Differences were considered statistically if the p-value was less than 0.05. To pre-determine sample sizes, we used G-power analysis based on pilot or previous studies. For animal studies, mice were assigned randomly, and sex was balanced in the various experimental groups; animals from the same litter were divided in different experimental groups; blinding of the investigator was applied to *in vivo* procedures and all data collection. Table S2 summarizes all the trials and read-outs performed.

## Results

### Setting up a method for SREBP2 delivery to striatal astrocytes in vivo

To over-express hSREBP2 in astrocytes in vivo in HD mice, we employed a recombinant serotype of an adeno-associated virus (AAV2/5) that is known to be highly specific for glial cells^24^. We modified the vector backbone to express the reporter gene *tdTomato* under the control a glial promoter, gfaABC1D^25^, a truncated variant of the human GFAP promoter (AAV2/5-gfaABC1D-tdTomato) or fused in-frame with the active N-terminal fragment of human SREBP2 (AAV2/5-gfaABC1D-hSREBP2-tdTomato). We first verified the spread and tropism over time of the AAV2/5-gfaABC1D-tdTomato vector using an in vivo method compatible with its application to R6/2 HD mice (Fig. 1A; Fig. S1A). Seven-week-old wild-type (wt) mice were administered a single unilateral intracranial injection of virus directly into the striatum. Four weeks later animals were sacrificed and subjected to different analyses, as summarized in Fig. 1B. Analysis of coronal sections of the brain by confocal microscopy demonstrated broad distribution of the AAV2/5-gfaABC1D-tdTomato in the infused striatum and in some parts of the cortex, indicating good spread from the injection site. In particular, tdTomato fluorescence was detectable over a span of 2.5 mm and it covered 61.51% ± 4.85 of the infused hemi-brain (Fig. 1C; Fig. S1B). AAV2/5-gfaABC1D-tdTomato tropism was analyzed by studying the co-localization of the tdTomato signal with the signals of neuronal (DARPP32, NeuN) and glial (GFAP, S100B, IBA1) markers. Figure 1D shows that AAV2/5 had a specific tropism for glial cells. In particular, the count of cells positive for tdTomato and for glial markers revealed that 80–90% of GFAP- or S100B-positive cells and about 40% of IBA1-positive cells also expressed tdTomato. In contrast, when counting cells positive for tdTomato and neuronal markers, no double-positive cells were found. A parallel study conducted by employing a recombinant Herpes Simplex Virus 1 (HSV1) vector as a backbone to deliver the same gfaABC1D-tdTomato construct (Fig. S1A) resulted in a widespread expression of the transgene throughout the injected striatum, although exclusively confined in neurons (Fig. S1B-C). We therefore proceeded with AAV2/5 to express hSREBP2 in astrocytes in HD mice.

**Figure 1.**
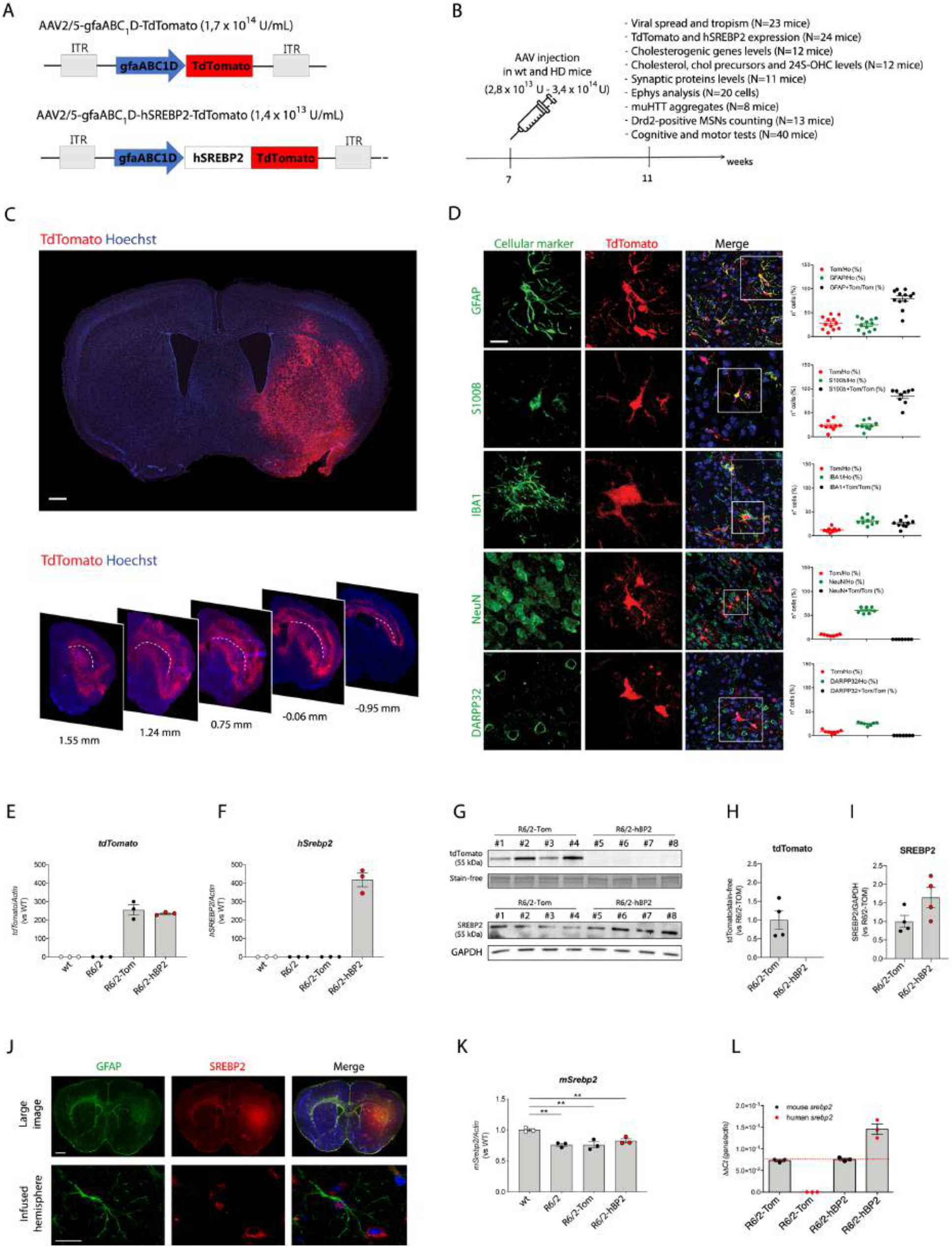
In vivo characterization of AAV2/5-gfaABC1D-tdTomato and AAV2/5-gfaABC1D-hSREBP2-tdTomato. **A–B.** Scheme of the experimental paradigm used in the study, and readouts performed. Mice at 7 weeks of age were infected in the right striatum with HSV1-JDNI8 or AAV2/5 expressing tdTomato under the control of the gfaABC1D promoter (A) and sacrificed 4 weeks later to study viral spread and tropism (B). **C.** Representative large images of coronal brain slices of wt mice infected with AAV2/5-gfaABC1D-tdTomato with immunostaining against tdTomato (red) to visualize AAV spread. Scale bar is 1000 *μ*m. **D.** Representative confocal images of coronal brain slices of wt mice infected with AAV2/5-gfaABC1D-TdTomato with immunostaining against tdTomato (red) and GFAP, IBA1, S100B, NeuN, or DARPP32 (green) to visualize viral tropism with relative quantification of the number of cells positive for tdTomato and for the specific markers; the number of cells positive for a specific marker is normalized on the number of the nuclei, counted in the field of view, and expressed as %). Scale bar is 5 *μ*m. **E–F.** mRNA levels of *tdTomato* (E) and *hSrebp2* (F) in the striatum and cortex from wt, R6/2, R6/2-Tom, and R6/2-hBP2 mice normalized on wt mice (*n* = 3 mice/group). **G–I.** Protein levels (G) and relative densitometry quantification of tdTomato (H) and hSREBP2 (I) in cytosolic fractions from the infused hemibrains from R6/2-Tom or R6/2-hBP2 (*n* = 4 mice/group). Stain-free imaging and GAPDH were used for normalization. **J.** Representative large image and high-magnification confocal image of coronal brain slices of R6/2-hBP2 mice with immunostaining against GFAP (green) and SREBP2 (red). **K.** mRNA levels of *mSrebp2* in the hemi-brains of wt and R6/2 mice, and in the infused hemi-brains of R6/2-Tom and R6/2-hBP2 mice (*n* = 3 mice/group). **L.** mRNA levels of *mSrebp2* and *hSrebp2* in the infused hemi-brains from R6/2-hBP2 mice (*n* = 3). Hoechst was used to counterstain nuclei (blue) in C, D, J. Scale bar is 1000 *μ*m (C), 5 *μ*m (D), 2000 *μ*m (J, up), and 10 *μ*m (J, down). Data (E, F, H, I, K, and L) are shown as scatterplot graphs with means ± SEM. Each dot corresponds to the value obtained from each animal. Statistics: one-way ANOVA with Newman–Keuls post-hoc test (\*\**p* < 0.01).

### Delivery of AAV2/5-gfaABC1D-hSREBP2-tdTomato to the HD striatum

To establish the efficiency of the *hSrebp2* transgene expression in vivo, AAV2/5-gfaABC1D-tdTomato or AAV2/5-gfaABC1D-hSREBP2-tdTomato were injected into the right striatum of 7 weeks old R6/2 mice (herein R6/2-Tom or R6/2-hBP2 respectively), a well-known HD animal model characterized by rapidly developing symptoms^26^. Four weeks later, when mice were at the end of the symptomatic stage, cortical and striatal tissues were collected and RNA transcripts and relative amounts of hSREBP2 protein were measured. For this experiment we decided to pool the cortex and striatum because in our preliminary analyses (Fig. 1C) some viral spread was also observed in the cortex.

As shown in Fig. 1E, RNA transcripts of *tdTomato* were not detected in the cortico-striatal samples of uninfected wt and R6/2 mice, while they were detected and quantified in R6/2-Tom and R6/2-hBP2 samples. As expected, RNA transcripts of *hSrebp2* were detected only in R6/2-hBP2 mice (Fig. 1F).

Western blotting revealed a tdTomato immunoreactive band in the lysates from all R6/2-Tom injected mice but not from the R6/2-hBP2 mice, indicating that *tdTomato* fused downstream to *hSrebp2* was not efficiently translated in AAV2/5-gfaABC1D-hSREBP2-tdTomato transduced cells (Fig. 1G and H). Importantly, western blot analysis with an antibody that recognizes both the endogenous (mouse) and exogenous (human) SREBP2 revealed an increase in total SREBP2 protein level in three out of four R6/2-hBP2 mice (Fig. 1I). The over-expression of hSREBP2 and its co-localization with astrocytes in the infused striatum was confirmed further in additional R6/2-hBP2 mice by immunofluorescent staining for GFAP and SREBP2 (Fig. 1J) and by performing stitching of multiple images following immunofluorescent staining that allowed a global view of hSREBP2 spread in the entire brain section (Fig. 1J).

As there are no antibodies available that reliably distinguish human from mouse SREBP2, we were not able to calculate the amount of the vector-derived hSREBP2 protein in the infected R6/2-hBP2 mice compared with the endogenous mouse SREBP2 (mSREBP2). To overcome this problem, we designed sets of primers to selectively quantify the absolute mRNA levels of *mSrebp2* and *hSrebp2* by qRT-PCR. Notably, we measured a significant reduction of the endogenous *mSREBP2* transcripts in R6/2, R6/2-Tom and R6/2-hBP2 mice compared with wt mice (Fig. 1K). Importantly, Fig. 1L shows that *hSrebp2* mRNA was between 1.7-fold and 2.1-fold more abundant than the endogenous transcript in R6/2-hBP2 mice. As these are whole tissue measurements, it is expected that mRNA level in the individually transduced cells may be higher.

### Glial hSREBP2 over-expression enhances cholesterol biosynthesis in HD mice

We next sought to test whether the AAV2/5-gfaABC1D-hSREBP2-tdTomato construct expresses a functional SREBP2 protein capable of increasing transcription of SREBP-controlled genes in the cholesterol biosynthesis pathway.

Gene expression analysis was performed on the pooled striatum and cortex from the injected hemisphere by measuring transcript levels of cholesterol biosynthesis genes (*Hmgcr*, *Mvk*, *Sqs/Fdft1*, *Cyp51, Dhcr7;* Fig. 2A) using qRT-PCR. The mRNA levels from all these genes were significantly reduced in R6/2 and R6/2-Tom mice compared with controls (Fig. 2B–F). Importantly, hSREBP2 over-expression in R6/2 mice rescued the transcript levels of *Hmgcr*, *Mvk*, *Fdft1*, and *Cyp51* (Fig. 2B-E) but not of *Dhcr7* (Fig. 2F). The transcript level of *Cyp46a1,* which encodes the enzyme that catalyzes the conversion of cholesterol to 24S-hydroxycholesterol (24S-OHC), was reduced in all R6/2 groups compared with controls but was not affected by exogenous hSREBP2 (Fig. 2G).

**Figure 2.**
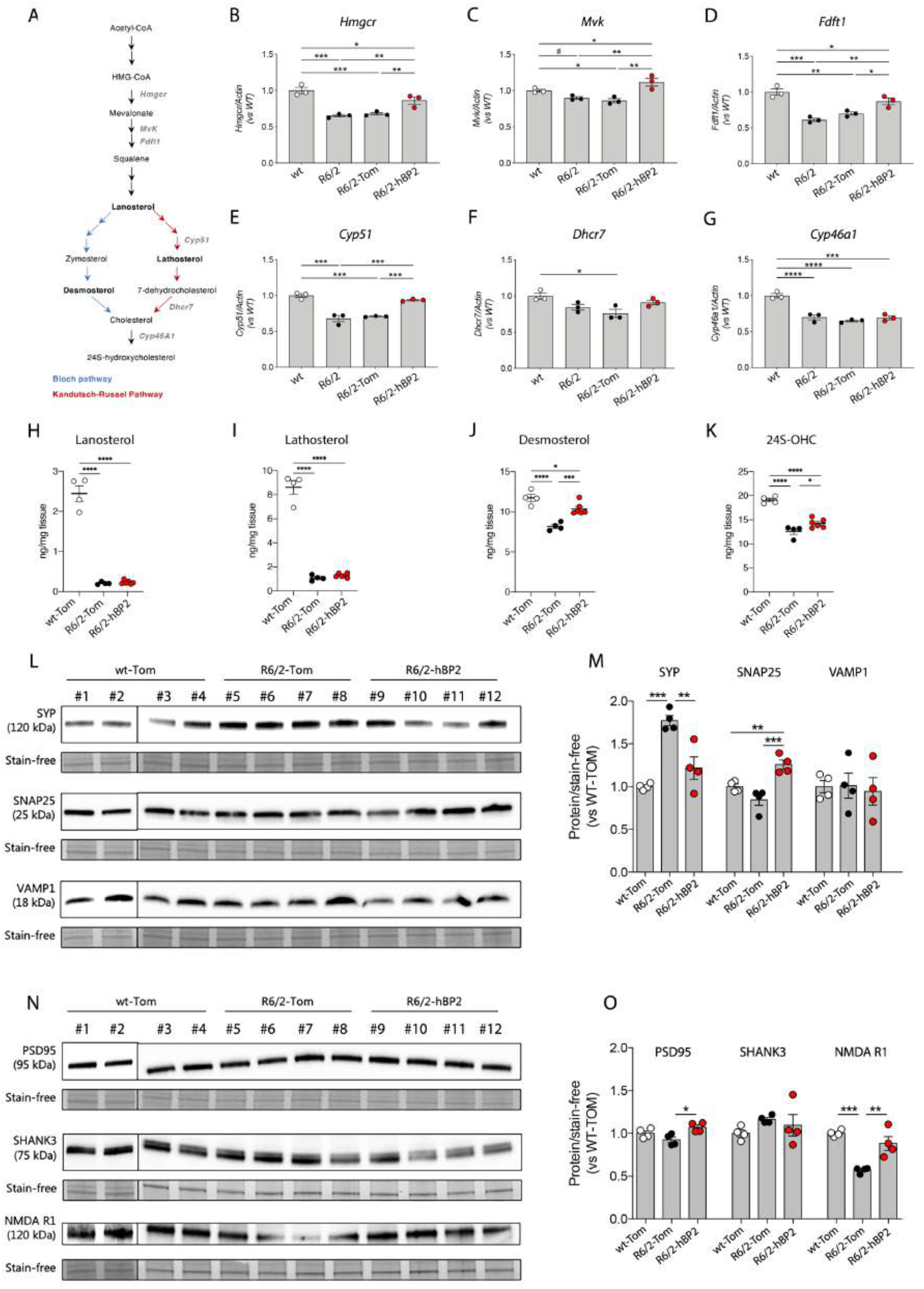
mRNA transcript levels of cholesterol biosynthesis genes and levels of synaptic proteins following glial hSREBP2 over-expression. **A.** Genes encoding enzymes of cholesterol biosynthesis whose expression was analyzed by qPCR (red). Products of the Bloch and Kandutsch-Russell pathways are in blue and pink, respectively. **B–G.** mRNA levels of hydroxymethylglutaryl-coenzyme A reductase (*Hmgcr*) (B), mevalonate kinase (*Mvk*) (C), squalene synthase/farnesyl-diphosphate farnesyl transferase 1 (*Sqs/Fdft1*) (D), cytochrome p450 lanosterol 14-alpha-demethylase (*Cyp51*) (E), 7-dehydroxycholesterol reductase (*Dhcr7*) (F), and cholesterol 24-hydroxylase (*Cyp46a1*) in the hemibrain from wt, R6/2, R6/2-Tom mice, and R6/2-hBP2 mice (*n* = 3 mice/group). **H–K.** Lanosterol (H), Lathosterol (I), desmosterol (J), and 24-OHC (K) levels measured by GC-MS in the infused striata of wt-Tom, R6/2-Tom, and R6/2-hBP2 mice (*n* = 4–6 mice/group). **L–M.** Protein levels (L) and relative densitometry quantification (M) of synaptophysin (SYP), synaptosome-associated protein 25 (SNAP25), and vesicle-associated membrane protein 1 (VAMP1) in synaptosomes purified from the infused hemi-brains from wt-Tom mice, R6/2-Tom mice, and R6/2-hBP2 mice (*n* = 4 mice/group). **N–O.** Protein levels (N) and relative densitometry quantification (O) of postsynaptic density protein 95 (PSD95), SH3 and multiple ankyrin repeat domains 3 (SHANK3), and N-methyl-D-aspartate receptor (NMDAR1) in synaptosomes purified from the infused hemi-brains from wt-Tom mice, R6/2-Tom mice, and R6/2-hBP2 mice (*n* = 4 mice/group). Stain-free imaging was used as a loading control and for normalization. Data (B-K, M, and O) are shown as scatterplot graphs with means ± SEM. Each dot corresponds to the value obtained from each animal. See Supplementary Figures S4 and S5 for full-length pictures of the blots shown in L and N. Statistics: one-way ANOVA with Newman–Keuls post-hoc test (\**p* < 0.05; \*\**p* < 0.01; \*\*\**p* < 0.001) or unpaired Student’s t-test (#*p* < 0.05).

To explore whether the increased transcription of SREBP2-controlled genes corresponded to an increased activity of the pathway, we measured cholesterol precursors (lanosterol, lathosterol, and desmosterol) as surrogate markers of cholesterol biosynthesis. As expected, the steady-state levels of all cholesterol precursors were reduced in the striata of R6/2-Tom mice compared with wt-Tom mice (Fig. 2H–J)^19,20,27,28^. hSREBP2 delivery in glial cells, although unable to affect lanosterol and lathosterol levels (Fig. 2H-I), led to a significant increase in desmosterol in the striata of R6/2-hBP2 mice compared with R6/2-Tom mice (Fig. 2J), suggesting a preferential stimulation of the Bloch pathway of cholesterol biosynthesis which is more related to astrocytes^29^ (Fig. 2A). This was accompanied by an increase in 24S-OHC (Fig. 2K), consistent with the notion that cholesterol synthesis and catabolism are in balance in the HD brain ^21^.

To assess whether the enhancement of cholesterol biosynthesis genes following striatal injection of AAV2/5-gfaABC1D-hSREBP2-tdTomato was accompanied by changes in the expression levels of genes involved in cholesterol efflux (*Abca1*), transport (*ApoE)*, and uptake (*Lrp1*), we measured their mRNA levels in cortico-striatal tissues. *Abca1* mRNA levels were increased in R6/2, R6/2-Tom, and R6/2-hBP2 mice with respect to wt mice (Fig. S2A), while *ApoE* and *Lrp1* mRNA levels were similar in all conditions (Fig. S2B and C), indicating that hSREBP2 delivery in glial cells did not affect transcript levels of these genes. Similarly, AAV2/5-gfaABC1D-hSREBP2-tdTomato did not influence transcript levels of genes involved in astrocytic homeostasis. The *Glt1* mRNA level was similarly reduced in R6/2, R6/2-Tom, and R6/2-hBP2 mice compared with controls (Fig. S2D), while levels of *S100B* and *Kir4.1* were similar in all groups (Fig. S2E and F). The mRNA levels of synapse-related genes such as *Bdnf*, *Cplx2*, *Shank3*, *Homer*, and *Gap43,* which were altered in R6/2 and R6/2-Tom mice, were not influenced by the forced expression of hSREBP2 in glial cells (Fig. S2G and H). Transcript levels of genes related to pathways such as energy and mitochondrial metabolism and autophagy were also not affected by the over-expression of hSREBP2 (Fig. S3A–I).

### Glial hSREBP2 over-expression influences the levels of synaptic proteins in HD mice

Since cholesterol secreted from glial cells participates in synapse formation and maintenance and influences the intracellular distribution of proteins involved in the synaptic machinery^6^, we decided to purify synaptosomes from the infused hemibrains of wt-Tom, R6/2-Tom, and R6/2-hBP2 mice and perform semi-quantitative western blot analysis for pre- and post-synaptic proteins.

We found that R6/2-Tom mice exhibited higher expression levels of the pre-synaptic protein SYP than wt-Tom mice and that glial hSREBP2 over-expression completely reversed this parameter (Fig. 2L and M). SNAP25 levels were similar in wt-Tom and R6/2-Tom, while a 38% increase was found in R6/2-hBP2 tissues (Fig. 2L and M). VAMP1 levels were similar in all conditions (Fig. 2L and M). Regarding the post-synaptic site, our densitometric analyses of the immunoreactive bands indicated a weak but significant increase (*p* = 0.0138) in PSD95 level in R6/2-hBP2 mice compared with wt-Tom and R6/2-Tom animals (Fig. 2N and O), even though differences between wt-Tom and R6/2-Tom mice were not observed (Fig. 2I and K). Conversely, SHANK3 levels were similar in all the tested mice, including R6/2-hBP2 mice (Fig. 2N and O). Finally, R6/2-Tom mice exhibited reduced NMDAR1 protein levels compared with wt-Tom mice, and forced expression of hSREBP2 normalized its levels to that of wt-Tom mice (Fig. 2N and O).

We conclude that glial hSREBP2 over-expression influences and partially reverses altered levels of synaptic proteins in synaptosomes of HD mice.

### Glial hSREBP2 over-expression restores synaptic communication in HD mice

As R6/2 striatal MSNs exhibit altered synaptic transmission^30,31^, we performed electrophysiological studies to evaluate whether synaptic dysfunction was reversed in R6/2-hBP2 mice.

In these studies, biocytin was included in the patch pipette to post hoc recover the morphological details of the recorded neurons (Fig. 3A). We first analyzed the passive membrane properties of striatal MSNs from brain slices of wt-Tom and R6/2-hBP2 mice. For R6/2-hBP2 mice, we compared data obtained from MSNs of the contralateral and the infused hemispheres. The comparison between R6/2-hBP2 and wt-Tom mice highlighted the fact that the membrane capacitance (C_m_), which is proportional to cell size, was significantly lower in R6/2-hBP2 compared with wt-Tom MSNs, while it was similar between the contralateral and the infused hemispheres of R6/2-hBP2 mice (Fig. 3B). On the other hand, input resistance (R_in_), reflecting the number of ion channels expressed by the cell, was significantly higher in both the infused and the contralateral hemispheres of R6/2-hBP2 compared with wt-Tom mice, and unchanged in the infused striata of R6/2-hBP2 mice compared with the contralateral ones (Fig. 3C).

**Figure 3.**
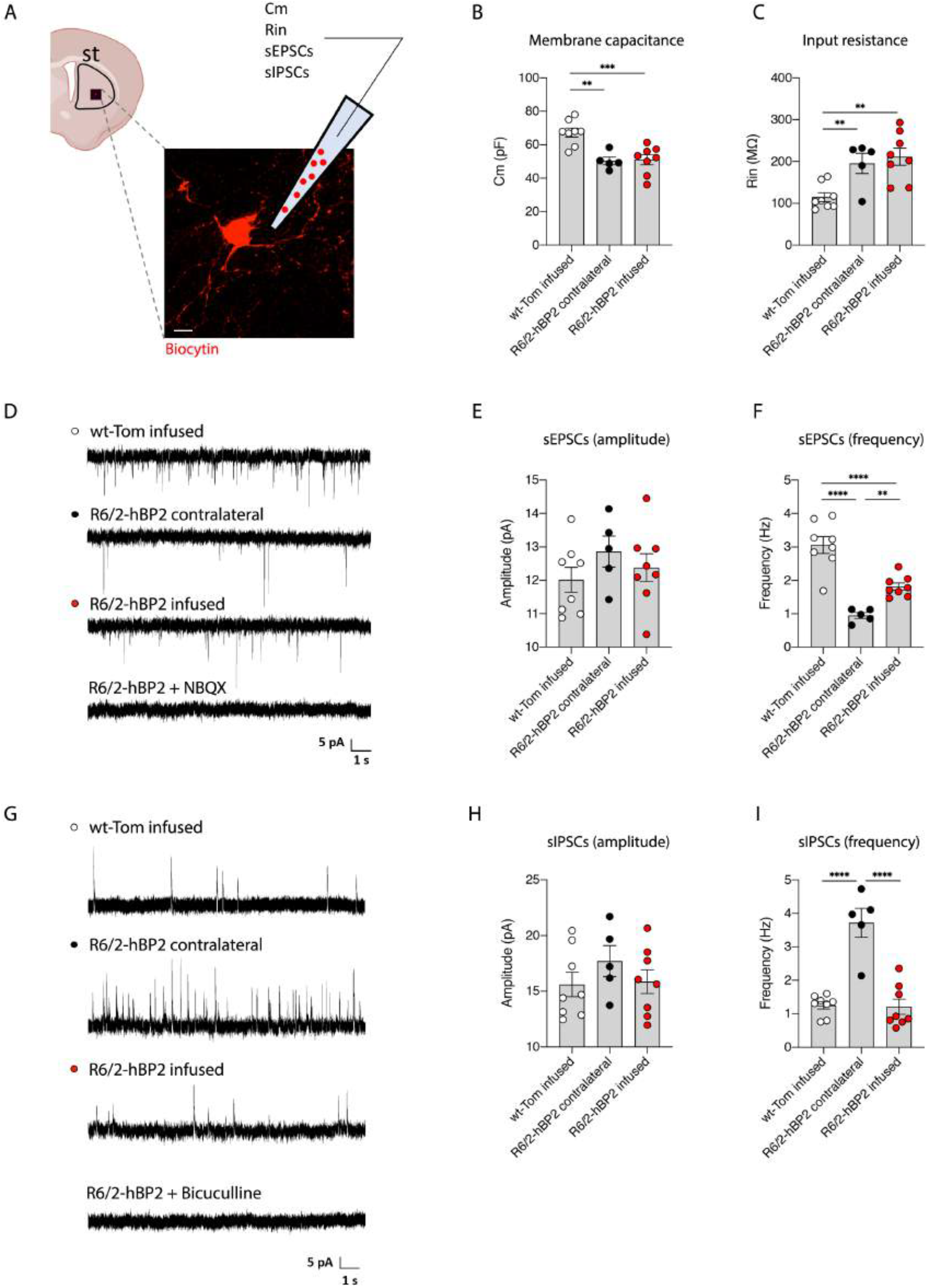
Electrophysiological analysis in MSNs of R6/2 mice following glial hSREBP2 over-expression. **A.** Schematic representation of the electrophysiological parameters analyzed in striatal MSNs of mice (stained with biocytin) following 4 weeks of AAV infusion. Scale bar is 5 *μ*m. **B–C.** Membrane capacitance (Cm, B) and input resistance (Rin, C) recorded from wt-Tom MSNs (n=8) and from the contralateral (*n* = 5) and infused (*n* = 8) MSNs of R6/2-hBP2 mice. **D–F.** Spontaneous EPSCs recorded from MSNs at a holding potential of −70 mV (D). Average amplitude (E) and average frequency (F) of EPSCs from wt and R6/2-hBP2 mice MSNs. **G–I.** Spontaneous IPSCs recorded from MSNs at a holding potential of 0 mV (G). Average amplitude (H) and average frequency (I) of IPSCs from wt and R6/2-hBP2 mice MSNs. Data (B, C, E, F, H, and I) are shown as scatterplot graphs with means ± SEM. Each dot corresponds to the value obtained from each animal. Statistics: one-way ANOVA with Newman–Keuls post-hoc test (\**p* < 0.05; \*\**p* < 0.01; \*\*\*\**p* < 0.0001; \*\*\**p* < 0.001).

To evaluate the effect of glial cholesterol on excitatory transmission, we recorded spontaneous excitatory postsynaptic currents (sEPSC) at a holding potential of −70 mV (Fig. 3D). The recorded events were AMPA-mediated since they were completely abolished by administration of 10 *μ*M NBQX (Fig. 3D). No significant differences in the average amplitude of sEPSC from MSNs were found between groups (Fig. 3E). In contrast, the average frequency of sEPSCs in MSNs from the contralateral hemisphere of R6/2-hBP2 mice was significantly lower compared with wt-Tom mice, and glial over-expression of hSREBP2 led to a partial rescue of this defect (Fig. 3F).

To test if the inhibitory synapses were also influenced by hSREBP2 transduction, we recorded spontaneous inhibitory synaptic currents (sIPSCs) at a holding potential of 0 mV (Fig. 3G). The recorded events were GABA-mediated since they were completely abolished by administration of 10 μM bicuculline (Fig. 3G). The average amplitude of sIPSCs was not affected (Fig. 3H). In contrast, the average frequency of sIPSCs in MSNs in the R6/2-hBP2-contralateral hemisphere was significantly higher compared with wt-Tom mice, and glial over-expression of hSREBP2 led to a normalization in the average frequency of sIPSCs in the infused hemisphere (Fig. 3I), bringing it closer to that in the wt-Tom mice.

### Glial hSREBP2 over-expression reduces muHTT aggregates in HD mice

A hallmark of HD is the presence of intracellular aggregates of muHTT^32^. To test whether glial over-expression of hSREBP2 in the HD striatum influenced muHTT aggregation, we performed immunofluorescence staining on coronal sections of brains from R6/2-Tom and R6/2-hBP2 mice by using the EM48 antibody, which is specific for the expanded polyQ tract prone to aggregate (Fig. 4A). As expected, tdTomato did not influence the number and the size of muHTT aggregates. Of note, the number and the size of EM48-positive aggregates were reduced in the infused striata of R6/2-hBP2 mice compared with the contralateral ones (Fig. 4B and C).

**Figure 4.**
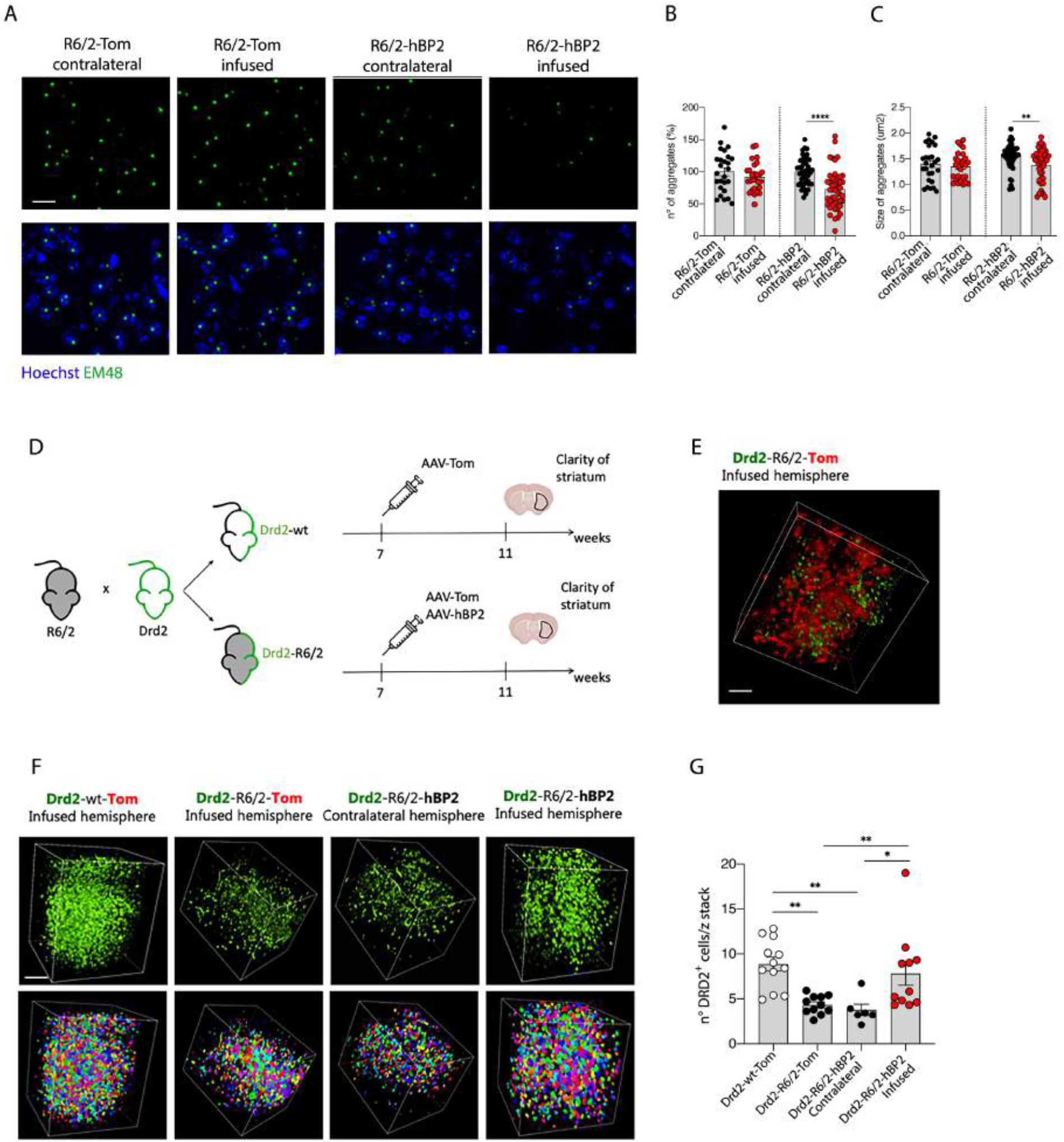
muHTT aggregation and number of Drd2 neurons in R6/2 mice following glial hSREBP2 over-expression. **A–C.** Immunolabeling of muHTT aggregates positive for EM48 antibody (green) (A) and relative quantification of number (B) and size (C) in infused and contralateral striata of R6/2-Tom mice or R6/2-hBP2 mice (*n* = 3–5/group). The number of aggregates in the infused hemisphere was normalized on the contralateral one. Hoechst (blue) was used to counterstain nuclei. Scale bar is 10 *μ*m. **D.** Experimental paradigm used in the CLARITY experiment. R6/2 mice were crossed with mice having Drd2-expressing MSNs tagged with GFP to obtain an HD line with neurons from the indirect pathway expressing GFP. Drd2 mice at 7 weeks of age were infected with AAV2/5-gfaABC1D-tdTomato (Drd2-wt-Tom) while Drd2-R6/2 mice were infected with AAV2/5-gfaABC1D-tdTomato (Drd2-R6/2-Tom) or with AAV2/5-gfaABC1D-hSREBP2-tdTomato (Drd2-R6/2-hBP2). Mice were sacrificed 4 weeks later and two 1 mm-thick brain coronal slices (comprehending the striatum) were prepared from each animal. From each slice, the portion including the infused and the contralateral striatum was isolated and clarified using the CLARITY technology (*n* = 5 mice/group). **E.** Representative two-photon imaging of the endogenous signals of GFP (green) and TdTomato (red) of 1-mm thick brain coronal slices from Drd2-wt-Tom (infused hemisphere). Scale bar is 100 *μ*m. **F–G.** Representative two-photon imaging (F, up) of the endogenous signal of GFP (green) of 1-mm thick brain coronal slices from Drd2-wt-Tom (infused hemisphere), Drd2-R6/2-Tom (infused hemisphere), and Drd2-R6/2-hBP2 (contralateral and infused hemisphere) with relative 3D reconstruction (F, down) and quantification (G). Scale bar is 200 *μ*m. Data (B, C and G) are shown as scatterplot graphs with means ± SEM. Each dot corresponds to an image. The number of neurons was normalized on the z-stack acquired. Statistics: one-way ANOVA with Newman–Keuls post-hoc test (\**p* < 0.05; \*\**p* < 0.01; \*\*\*\**p* < 0.0001).

### Glial hSREBP2 over-expression increases the number of Drd2-expressing MSNs in HD mice as detected by CLARITY technology

Neuronal dysfunction of striatal neurons precedes their degeneration, and the dopamine receptor D2 (DRD2) MSNs of the indirect pathway are the first affected in HD models and patients^33,34,35,36^. Although R6/2 mice (similar to other HD mouse models) do not show overt signs of neuronal loss^26^, the downregulation of *Drd2* transcript in striatal MSNs is an established early mark of disease progression that has been proposed as a sensitive measure of the effects of therapeutics^37^. In particular, Crook and Housman developed a Drd2 dysregulation assay by crossing Drd2-eGFP mice with different HD mice to obtain animals with DRD2-MSNs labeled with eGFP, and then used FACS sorting as a quantitative and reproducible readout of the efficacy of their therapeutics^37^.

Here we adopted the same strategy to obtain animals with DRD2-MSNs expressing eGFP (herein Drd2-wt and Drd2-R6/2 mice) but used the CLARITY, a tissue clearing technology that allows to obtain an accurate measurement of the total number of MSNs expressing DRD2 in a large volume of the vector-injected striatum (approx. 108±5 *μ*m^3^), without dissociation.

To accomplish this goal, we injected AAV2/5-gfaABC1D-tdTomato or AAV2/5-gfaABC1D-hSREBP2-tdTomato into the right striata of 7-week-old Drd2-wt and Drd2-R6/2 mice (herein Drd2-wt-Tom, Drd2-R6/2-Tom, and Drd2-R6/2-hBP2). Animals were sacrificed 4 weeks later. From each animal, two brain coronal slices which were 1 mm thick (including the striatum) were prepared from the infused and contralateral hemispheres and the area corresponding to the striatum was isolated and clarified using the CLARITY technology (Fig. 4D). The images acquired with two-photon microscopy in the clarified striata of Drd2-wt-Tom mice confirmed distinct signals of tdTomato and eGFP, which labeled infected glial cells and DRD2-MSNs, respectively (Fig. 4E). The subsequent quantification of the eGFP signal in the different experimental groups revealed a reduced number of DRD2-MSNs in the infused striata of Drd2-R6/2-Tom mice and in the contralateral striata of Drd2-R6/2-hBP2 compared with Drd2-wt-Tom mice (Fig. 4F and G). Of note, the number of DRD2-MSNs measured in the infused striata of Drd2-R6/2-hBP2 was similar to that quantified in Drd2-wt-Tom mice (Fig. 4F and G), indicating that over-expression of hSREBP2 in astrocytes allows these neurons to regain DRD2 expression and, supposedly, functionality.

### Glial hSREBP2 over-expression ameliorates motor defects and completely restores cognitive decline in HD mice

To assess whether the described amelioration of synaptic activity and disease phenotypes following over-expression of hSREBP2 in astrocytes was correlated with a beneficial effect on behavioral abnormalities, we subjected wt-Tom, R6/2-Tom, and R6/2-hBP2 mice to motor and cognitive tasks. As expected, 11-week-old R6/2-Tom mice exhibited a severe hypokinetic phenotype as demonstrated by reduced global activity, total distance traveled, and number of rearings in the activity cage test (Fig. 5A–C). Glial over-expression of hSREBP2 led to a slight but significant rescue of these parameters (Fig. 5A–C). We also evaluated the time that mice spent exploring the peripheral or central area of the arena during the activity cage test as a measure of anxiety-like behavior (Fig. 5D). R6/2-Tom mice showed anxiety-related behavior as they spent more time in the periphery compared with wt-Tom mice. Of note, R6/2-hBP2 mice behaved similarly to wt-Tom mice (Fig. 5E). When we measured the neuromuscular function of mice with the grip-strength test, we observed reduced muscle strength in R6/2-Tom mice compared with wt-Tom mice and, again, glial over-expression of hSREBP2 had a beneficial effect on this feature (Fig. 5F). Furthermore, we performed the paw-clasping test to study the clasping behavior of mice as an index of neuronal dysfunction. As expected, R6/2-Tom mice had a worse performance compared with wt-Tom, but in this case the treatment was not able to significantly rescue this behavior (Fig. 5G). Finally, to analyze the effect of treatment on cognitive decline, we performed the novel-object recognition (NOR) test: at 11 weeks of age, R6/2-Tom mice were not able to discriminate between the novel object and the familiar one, in contrast to the wt-Tom mice. Of note, glial over-expression of hSREBP2 completely prevented cognitive decline and R6/2-hBP2 mice behaved as the wt-Tom (Fig. 5H).

**Figure 5.**
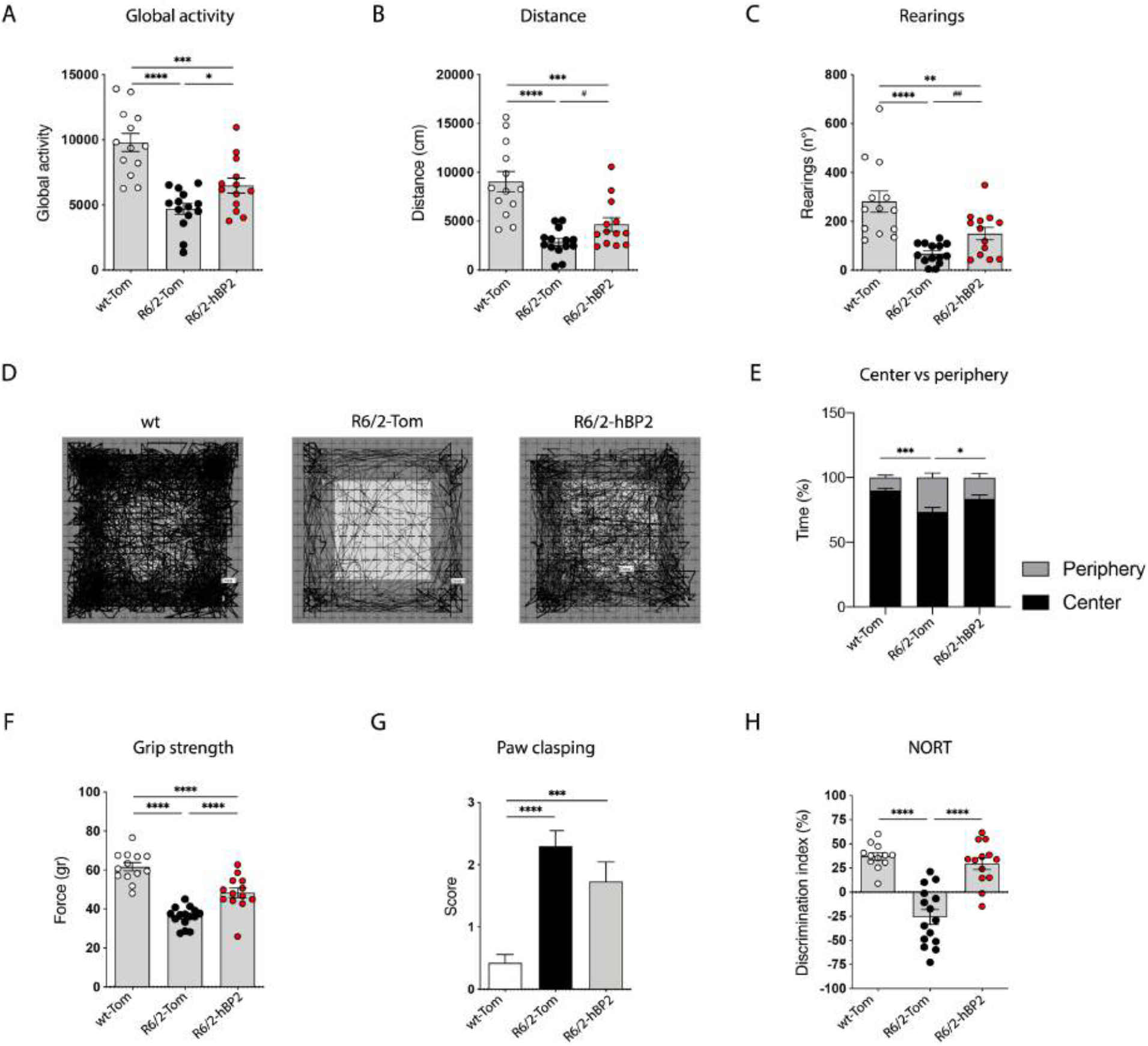
Cognitive and motor abilities of R6/2 mice following glial hSREBP2 over-expression. **A–E.** Global activity (A), distance traveled (B), and number of rearings (C) in an open-field test in wt-Tom (*n* = 13), R6/2-Tom (*n* = 14), and R6/2-hBP2 (*n* = 13). Representative track plots (D) from the open-field test from wt-Tom (*n* = 13), R6/2-Tom (*n* = 14), and R6/2-hBP2 (*n* = 13) and relative quantification (E) of the time spent (%) in the center and in the periphery of the arena. **F.** Grip strength (grams) in wt-Tom (*n* = 13), R6/2-Tom (*n* = 15), and R6/2-hBP2 (*n* = 13). **G.** Paw clasping in wt-Tom (*n* = 13), R6/2-Tom (*n* = 15), and R6/2-hBP2 (*n* = 13). **H.** Discrimination index (DI %) in the novel object recognition test of wt-Tom (*n* = 12), R6/2-Tom (*n* = 15), and R6/2-hBP2 (*n* = 13). DI above zero indicates a preference for the novel object; DI below zero indicates a preference for the familiar object. Data (A-C and F-H) are shown as scatterplot graphs with means ± SEM. Each dot corresponds to the value obtained from each animal. Statistics: one-way ANOVA with Newman–Keuls post-hoc test (\**p* < 0.05; \*\**p* < 0.01; \*\*\**p* < 0.001; \*\*\*\**p* < 0.0001) or unpaired Student’s t-test (#*p* < 0.05; ##*p* < 0.01).

## Discussion

In this work, we highlight the therapeutic value of forcing cholesterol biosynthesis in vivo in an HD mouse model via a gene therapy approach that delivers the active form of the master transcription factor SREBP2 specifically to astrocytes. In the brain, the physiological cross-talk between astrocytes and neurons provides the latter with the cholesterol produced by the former, which is then utilized by neurons to support their function. This cross-talk is severely damaged in vitro in the presence of muHTT, due to reduced activity of SREBP2, but it can be rescued when HD astrocytes over-expressing SREBP2 are co-cultured with HD neurons. In these conditions, we showed that neuronal function and morphology are fully reestablished^22^. Here, we now provide in vivo evidence that this altered communication between astrocytes and neurons is a key pathological feature of HD astrocytes, which can be targeted at a cellular level and their pathology reversed, leading to substantial amelioration of HD phenotypes in the living animals. Accordingly, we show that hSREBP2 delivery to astrocytes in vivo, by enhancing the cholesterol biosynthesis pathway, can revert molecular, biochemical, electrophysiological, and behavioral abnormalities associated with neuronal dysfunction typically observed in the R6/2 mouse model.

We show that unilateral striatal AAV-delivery of SREBP2 in glial cells is sufficient to completely rescue cognitive decline in R6/2 mice. Similar data were obtained upon cholesterol delivery to the brain of this mouse model via brain-permeable polymeric nanoparticles or mini-pumps^27,28^. The fact that genetic deletion of SCAP or SREBP2 in astrocytes of adult mice results in cognitive defects^38,39^ confirms the role of astrocytes in memory and cognition in vivo, and that altered cross-talk between astrocytes and neurons at synaptic contacts may contribute to cognitive disturbances^40^. Moreover, cholesterol synthesis is downregulated in aging astrocytes^41,42^, suggesting that lack of cholesterol production in these cells may contribute to the synaptic and cognitive defects observed in aging. Here, we provide further evidence that cholesterol produced from astrocytes in vivo is also a critical player in memory recognition in the HD context.

Unilateral AAV-delivery of SREBP2 also ameliorated motor defects in R6/2 mice, suggesting that this approach is able to produce the optimal content of newly synthetized cholesterol needed to partially restore motor-dependent circuits. These results are in agreement with our recent work that evaluated the effect of three escalating doses of cholesterol infused directly into the striata of R6/2 mice through osmotic mini-pumps. Indeed, we demonstrated that all three doses prevented cognitive defects, while only the highest dose attenuated motor phenotypes^28^.

Mechanistically, our findings indicate that hSREBP2 over-expression in glial cells carries out its beneficial effect by acting on different aspects of the disease. The synaptic effect of AAV delivery of hSREBP2 is certainly the relevant one, and it is supported by the evidence of normalization in the level of synaptic proteins and of both inhibitory and excitatory synaptic communication of MSNs. Newly synthetized cholesterol from astrocytes may facilitate synaptic vesicle fusion and exocytosis^43^ or bind to proteins involved in synaptic transmission^8,44^.

Our current data also suggest that the mechanisms by which stimulation of cholesterol biosynthesis counteracts synaptic dysfunction in HD mice do not involve major transcriptional changes since mRNA levels of key synaptic genes, which are known to be altered in HD, are unaffected by hSREBP2 delivery. Nor are the mRNA levels of a subset of genes involved in glial or energy metabolism influenced by exogenous hSREBP2.

At the neuropathological level, increased hSREBP2 in the striatum of R6/2 mice reduced the number and size of muHTT aggregates, supporting the notion that enhanced cholesterol biosynthesis may participate in inducing clearance pathways, as we recently showed in the striata of HD mice following striatal infusion of cholesterol through osmotic mini-pumps^28^. The fact that in the current study the transgene was delivered to astrocytes suggests that clearance may also occur by a non-cell autonomous mechanism.

Importantly, hSREBP2 over-expression in glial cells rescues specific neurochemical features in MSNs. In fact, reduced striatal DRD2 was consistently demonstrated in patients^45,46,47,48,49^ and murine models^27,35,50,51^ and its dysregulation is a sensitive measure of HD pathology^37^. Of note, a loss of dopamine receptors has also been correlated with early cognitive decline^52^. Our CLARITY approach applied to the striatum allowed for the first time an accurate measurement and quantification of striatal MSNs in situ and confirmed the decrease in the level of DRD2 in MSNs^37^. Importantly, we show that HD neurons, which receive more cholesterol from hSREBP2 over-expressing astrocytes, increase DRD2 expression and are quantifiable through two-photon imaging. These findings also suggest that within the striatum, the astrocyte-neuron signaling may be synapse-and cell-specific by acting at least on the DRD2-MSNs, as recently proposed^53^. The observed rescue in DRD2-MSNs, and in other synaptic features, may be favored by a closer contact between neurons and astrocytes, which is reduced in R6/2 astrocytes^54^.

In a similar gene therapy approach, the gene encoding for CYP46A1, the enzyme that converts cholesterol into the neuronal cholesterol catabolite 24S-OHC, was delivered into striatal neurons of HD mice by use of an AAVrh10 vector^55,56^. The treatment decreased the number and size of muHTT aggregates and improved motor deficits in R6/2 mice^54^ and provided long-lasting improvement in zQ175 HD mice^56^. In both studies, levels of cholesterol precursors were restored or increased in the HD striatum following AAV-delivery of CYP46A1. In our study, we show increased levels of 24S-OHC following AAV-delivery of hSREBP2 in glial cells, although *Cyp46A1* mRNA remained unchanged, supporting the close balance between cholesterol synthesis and catabolism in the diseased brain.

In conclusion, our findings confirm the role of astrocytes as important regulatory cell types in brain function and behavior^40,57^ and in neurodegenerative disorders, including HD^58,59^. By targeting astrocytic cholesterol signaling in the striatum we were able to rescue biochemical, neuropathological, functional, and behavioral features in HD mice. Thus, SREBP2 gene transfer to astrocytes may be a therapeutic option to fight core features of this disease.

## Acknowledgments

The authors acknowledge Dr. Cesare Covino and Advanced Light and Electron Microscopy BioImaging Center (ALEMBIC) at San Raffaele Institute, Milan (EuBI_PIRI081); Dr. Laura Madaschi and NOLIMITS, an advanced imaging facility established by the University of Milan; Monica Favagrossa and Marta Vittani for technical contribution on this study during the preparation of their experimental thesis.

This research was partially supported by Telethon Foundation (GGP17102), NeurostemcellRepair (FP7, GA no. 602278, 2013-17), NSC-Reconstruct (H2020, GA no 874758, 2020-23) (to E.C.); by Linea 2-2017, Department of Biosciences, University of Milan (to M.V.); by the Italian Ministry of Education, University and Research (MIUR): Dipartimenti di Eccellenza Program (2018–2022) - Dept. of Biology and Biotechnology “L. Spallanzani”, University of Pavia (to G. Biella, F.T., M.C.). F.T. was supported by Fondazione Umberto Veronesi.

## Author contributions

M.V., M.S. and E.C. conceived the study; G.V. and M.S. provided HSV vectors; L.Z. and M.G. provided AAV vectors; G. Birolini and M.V. performed *in vivo* experiments, including mice infection with viral vectors and behavioral analysis; G. Birolini performed immunostaining experiments and provided confocal images and quantification; G. Birolini, M.V. and G.V. performed Clarity experiments; G. Birolini performed western blot analysis; P.C. provided advices for qRT-PCR experiments; C. Cordiglieri gave scientific and technical assistance for the quantification of 2-phton images; M.V. produced the plasmids and contributed to perform qRT-PCR experiments; C.M., F.T., and G. Biella performed and analyzed the electrophysiological recordings; C.Caccia, V.L. and F.T. performed and analysed mass spectrometry data; M.V. and G. Birolini collected study data and performed statistical analyses; M.V. and E.C. oversaw and coordinated responsibility for all research activities and their performance and provided experimental advice throughout the work; E.C. and M.V. secured the funding, the collaborations and the execution of the entire project. M.V., G. Birolini, and E.C. wrote the paper that has been edited and reviewed by all authors.

**Figure S1.**
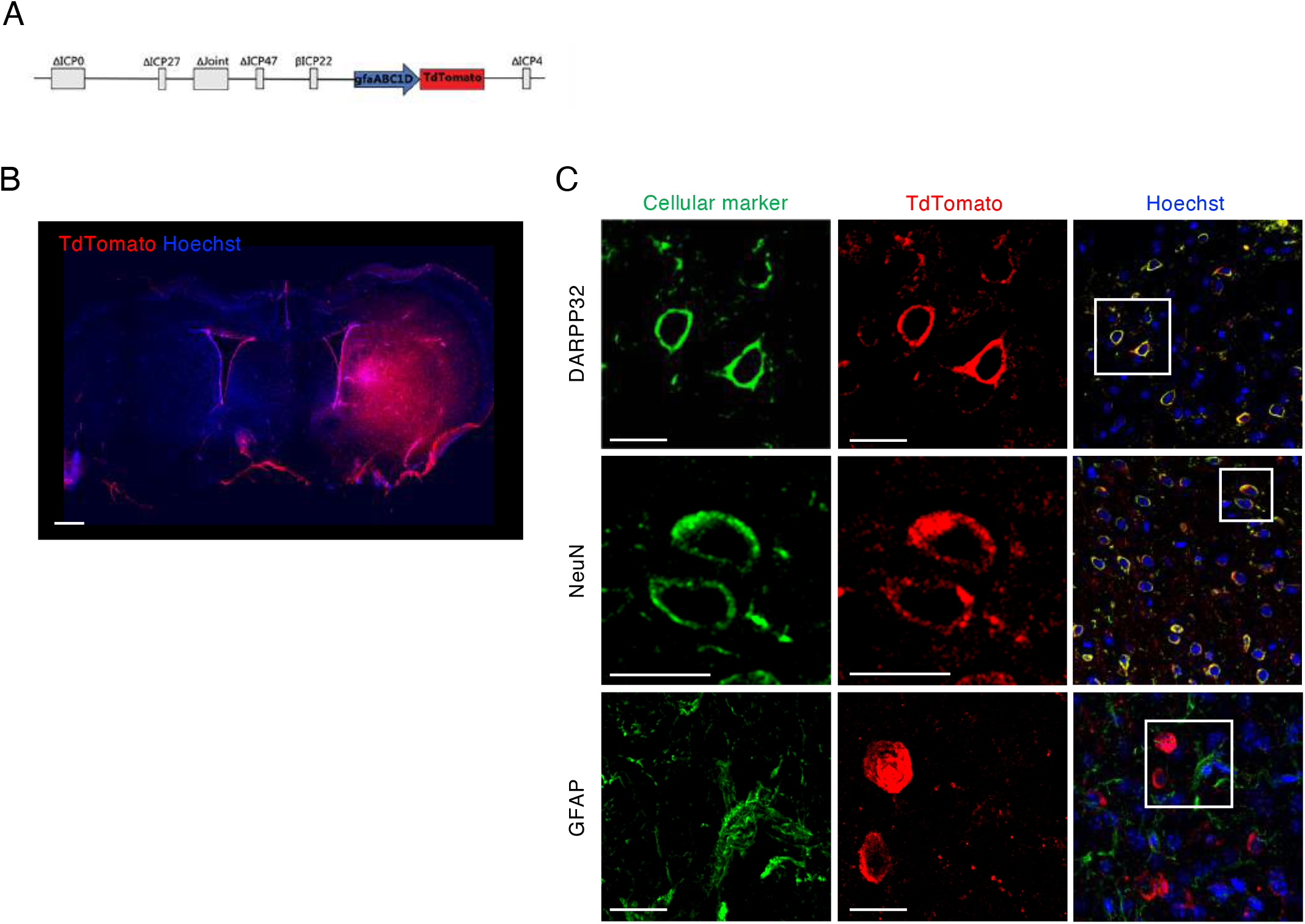
Spread and tropism of HSV1/JDNI8-gfaABC1D-TdTomato. **A.** Representative large images of coronal brain slices of wt mice infected with HSV/JDNI8-gfaABC1D-TdTomato with immunostaining against TdTomato (red) to visualize viral spread. Hoechst were used to counterstain nuclei (blue). Scale bar is 1000 *μ*m. **B.** Representative confocal images of coronal brain slices of wt-HSV/JDNI8-gfaABC1D-TdTomato mice with immunostaining against TdTomato (red) and DARPP32, NeuN, and GFAP (green) to visualize viral tropism. Hoechst were used to counterstain nuclei (blue). Scale bar is 10 *μ*m.

**Figure S2.**
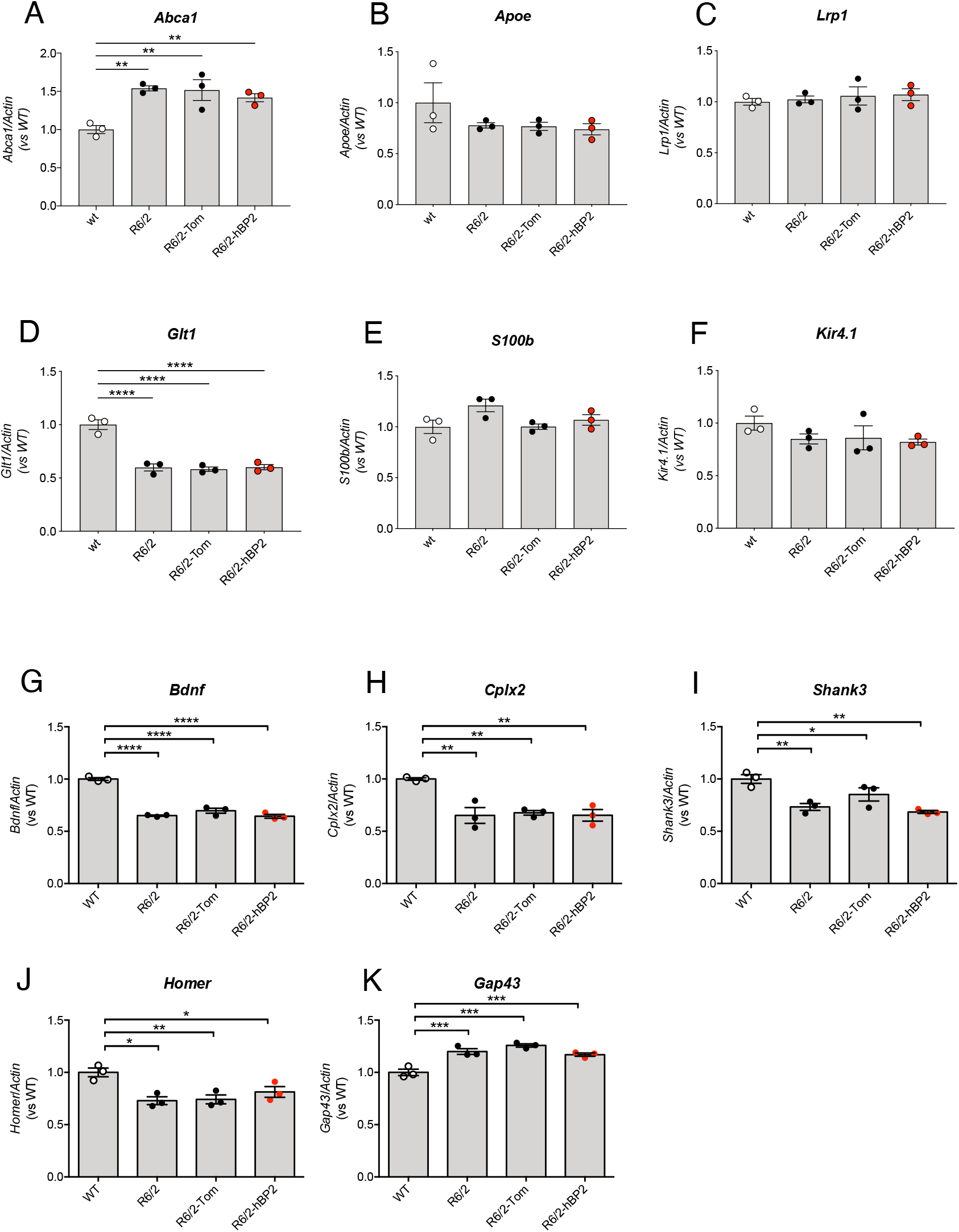
Gene expression analysis of genes involved in cholesterol metabolism, glial metabolism and synaptic activity. **A-D.** mRNA level of ATP-binding cassette transporter (*Abca1*) (A), Apolipoprotein E (*ApoE*) (B), Low density lipoprotein receptor-related protein 1 (*Lrp1*) (C), and Cholesterol 24-hydroxylase (*Cyp46a1*) (D). in the hemi-brain of wt and R6/2 mice and in the infused hemibrain from R6/2-Tom, and R6/2-hBP2 mice (*n* = 3 mice/group). **E-G.** mRNA level of Glutamate transporter 1 (*Glt1*) (E), S100 calcium binding protein B (*S100b*) (F), and potassium inwardly-rectifying channel, subfamily J, member 10 (*Kir4*.1) (G) in the hemi-brain of wt and R6/2 mice and in the infused hemibrain from R6/2-Tom, and R6/2-hBP2 mice (*n* = 3 mice/group). **H-L.** mRNA levels of Brain derived neurotrophic factor (*Bdnf*) (H), Complexin-2 (*Cplx2*) (I), SH3 and multiple ankyrin repeat domains 3 (*Shank3*) (J), Homer scaffolding protein (*Homer*) (K), and Growth-associated protein 43 (*Gap43*) (L) in the hemi-brain of wt and R6/2 mice and in the infused hemibrain from R6/2-Tom, and R6/2-hBP2 mice (*n* = 3 mice/group). Data (A-L) are shown as scatterplot graphs with means ± SEM. Each dot corresponds to the value obtained from each animal. Statistics: one-way ANOVA with Newman–Keuls post-hoc test (\**p*<0.05; \*\**p*<0.01; \*\*\**p*<0.001; \*\*\*\**p*<0.0001).

**Figure S3.**
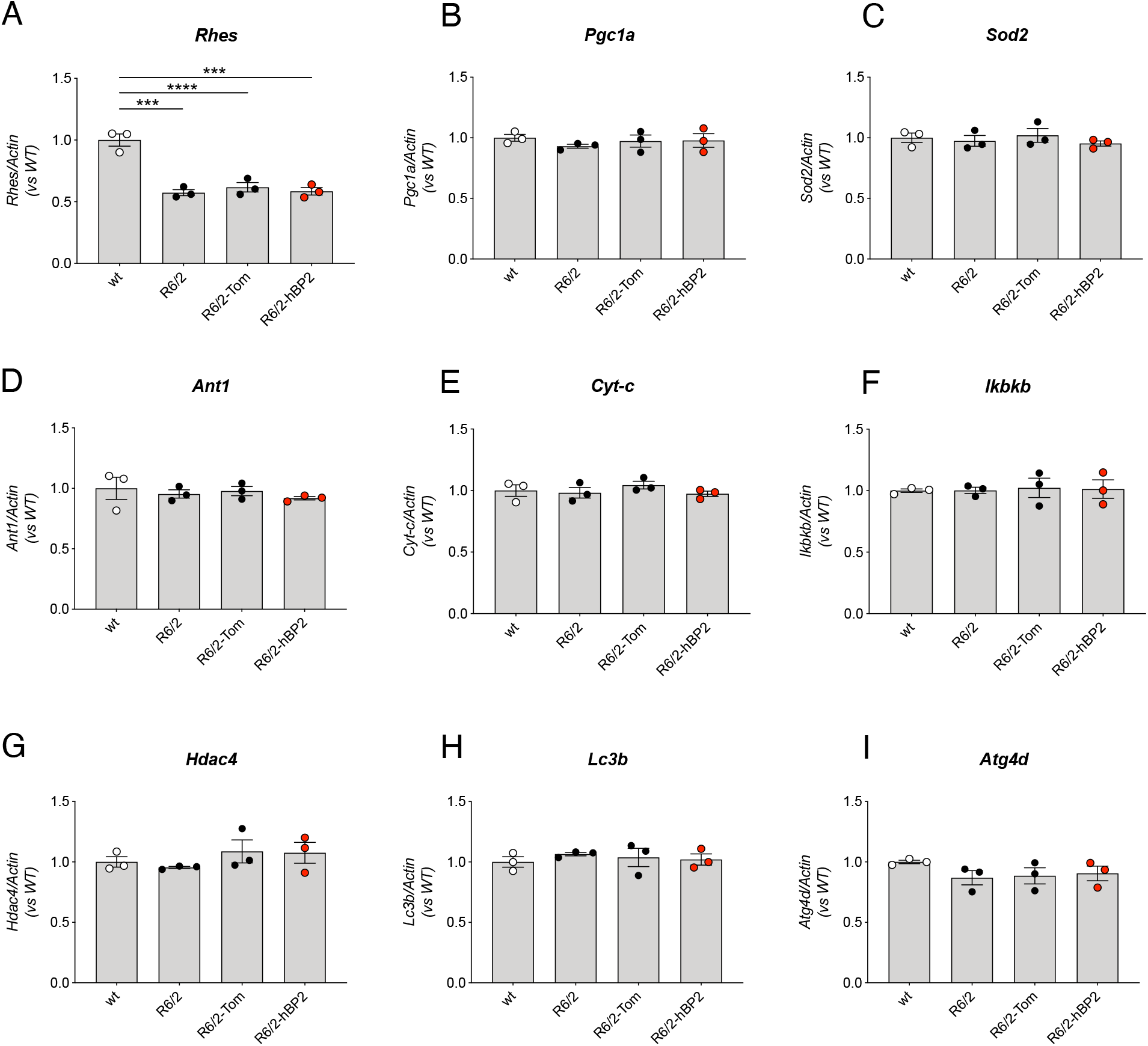
Gene expression analysis of genes involved in energy metabolism and autophagy. **A-I**. mRNA levels of GTP-binding protein Rhes (*Rhes*) (A), Peroxisome proliferator-activated receptor gamma coactivator 1-alpha (*Pgc1a*) (B), Superoxide dismutase 2 (*Sod2*) (C), Peroxisomal adenine nucleotide transporter 1 (*Ant1*) (D), cytochrome complex (*Cyt-c*) (E), Inhibitor Of Nuclear Factor Kappa B Kinase Subunit Beta (*Ikbkb*) (F), Histone deacetylase 4 (*Hdac4*) (G), Microtubule-associated proteins 1A/1B light chain 3B (*Lc3b*) (H), and Autophagy Related 4D Cysteine Peptidase (*Atg4d*) (I) in the hemi-brain of wt and R6/2 mice and in the infused hemibrain from R6/2-Tom, and R6/2-hBP2 mice (*n* = 3 mice/group). Data (A-I) are shown as scatterplot graphs with means ± SEM. Each dot corresponds to the value obtained from each animal. Statistics: one-way ANOVA with Newman–Keuls post-hoc test (\*\*\**p*<0.001; \*\*\*\**p*<0.001).

**Figure S4.**
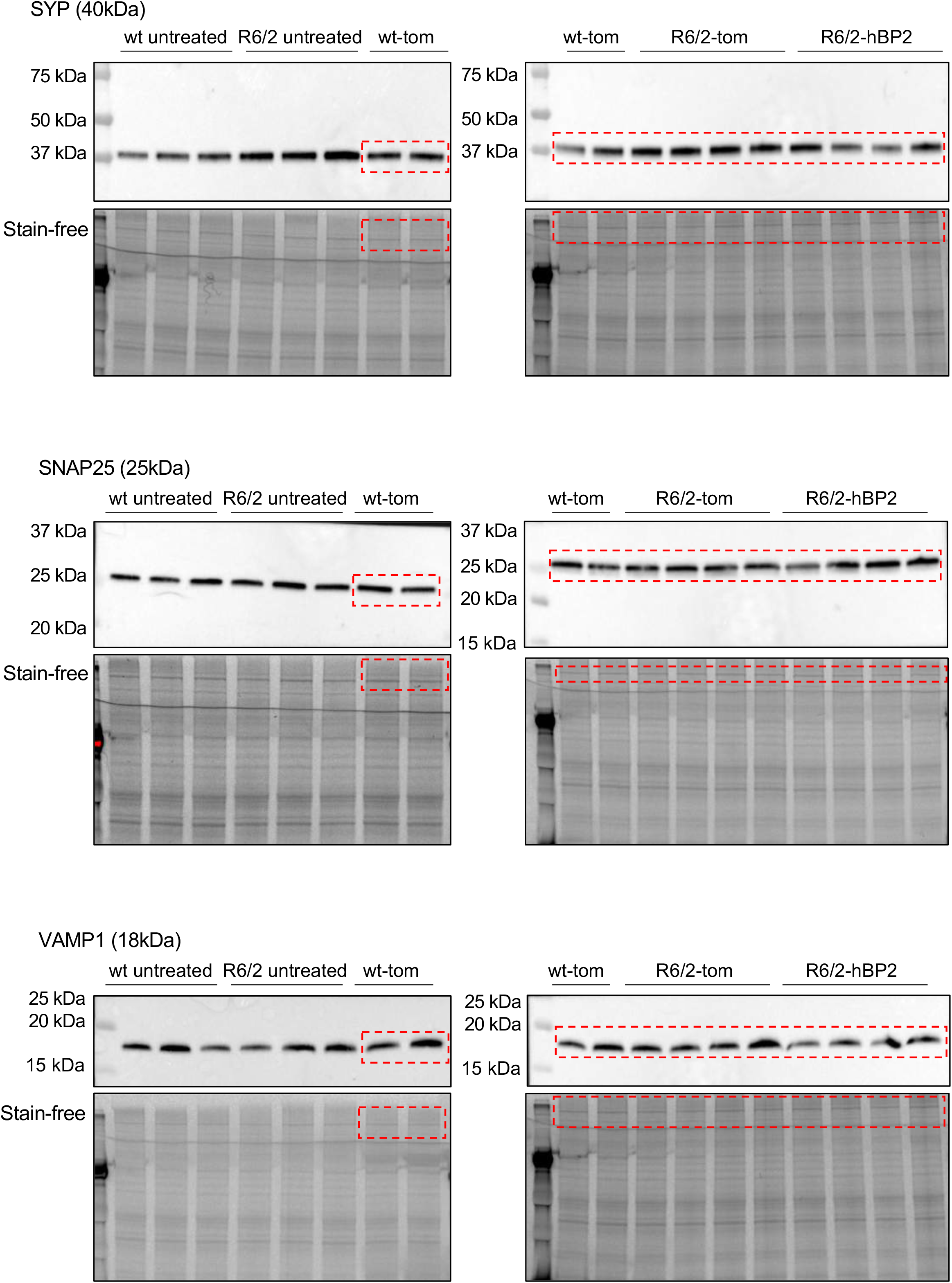
Source Data_Figure 2L. Full-length pictures of the plots presented in Figure 2L.

**Figure S5.**
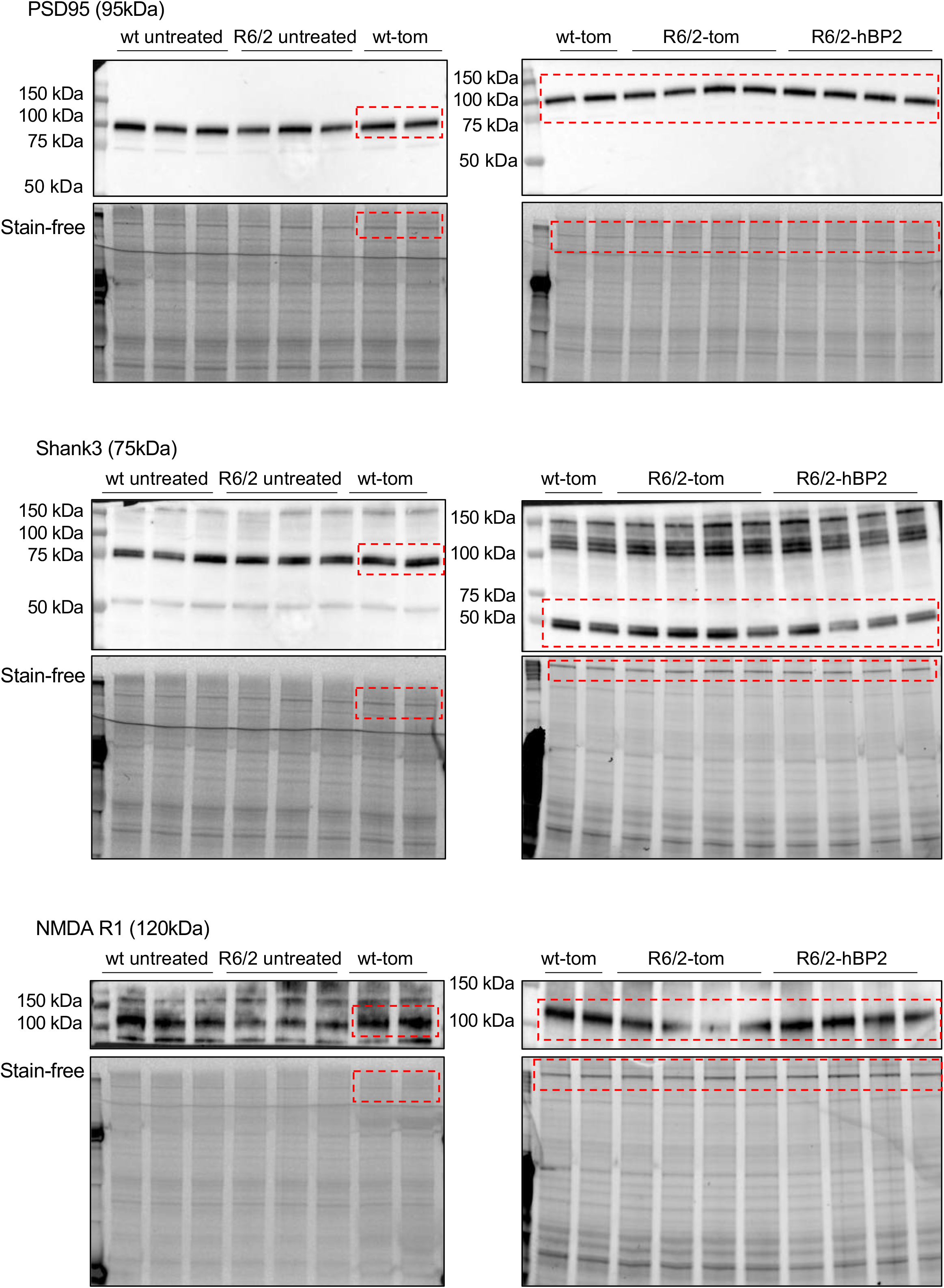
Source Data_Figure 2N. Full-length pictures of the plots presented in Figure 2N.

**Table S1.**
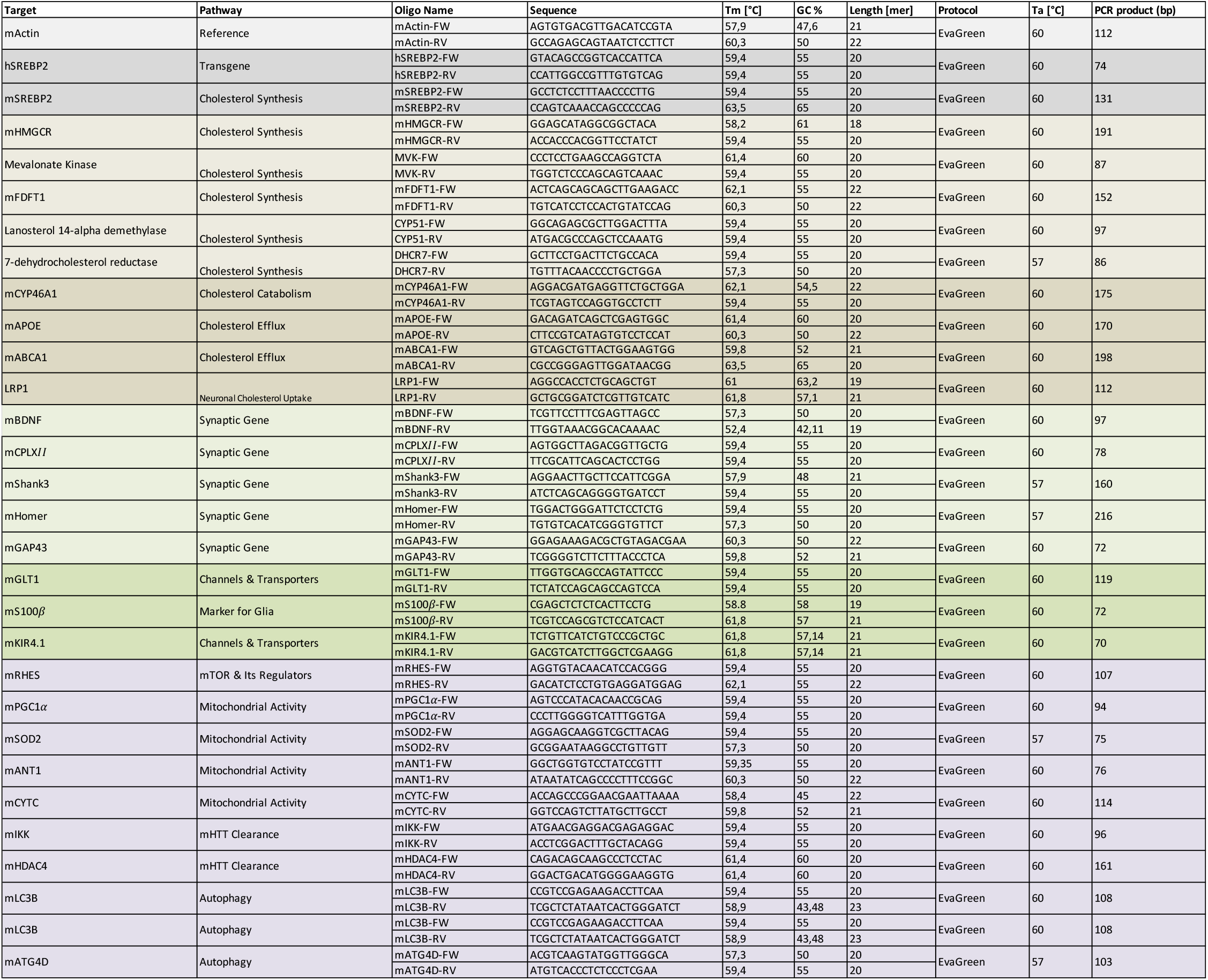
Primers List.

**Table S2.** Animals used in all the experiments.

| Table S2. Animals used in all the experiments |  |  |  |  |  |  |  |  |  |  |
| --- | --- | --- | --- | --- | --- | --- | --- | --- | --- | --- |
| N° Trial | Date | Experimental Groups | Spread and tropism | qRT-PCR | Synaptosomes | Ephys analysis | Drd2 quantification | muHTT analysis | Mass spect analysis | Behavioral analysis |
|  |  |  | N mice | N mice | N mice | N mice | N mice | N mice |  | N mice |
| 1 | Aug-Sept 2017 | wt-HSV | 4 |  |  |  |  |  |  |  |
|  |  | wt-Tom | 3 |  |  |  |  |  |  |  |
|  |  | R6/2-Tom | 3 |  |  |  |  |  |  |  |
|  |  | R6/2-hBP2 | 5 |  |  |  |  |  |  |  |
| 2 | Nov 2017 | wt |  | 3 |  |  |  |  |  |  |
|  |  | R6/2 |  | 3 |  |  |  |  |  |  |
|  |  | R6/2-Tom | 3 | 3 |  |  |  |  |  |  |
|  |  | R6/2-hBP2 | 2 | 3 |  |  |  |  |  |  |
| 3 | Ott 2018 | wt-Tom |  |  | 4 |  |  |  |  |  |
|  |  | R6/2-Tom |  |  | 4 |  |  |  |  |  |
|  |  | R6/2-hBP2 | 3 |  | 4 |  |  |  |  |  |
| 4 | Dic 2019 | wt |  |  |  | 5 |  |  |  |  |
|  |  | R6/2-hBP2 |  |  |  | 5 |  |  |  |  |
| 5 | Gen 2020 | wt-Tom |  |  |  |  |  |  |  | 8 |
|  |  | R6/2-Tom |  |  |  |  |  | 3 |  | 8 |
|  |  | R6/2-hBP2 |  |  |  |  |  | 5 |  | 7 |
| 6 | Feb 2020 | Drd2-wt-Tom |  |  |  |  | 5 |  |  |  |
|  |  | Drd2-R6/2-Tom |  |  |  |  | 4 |  |  |  |
|  |  | Drd2-R6/2-hBP2 |  |  |  |  | 4 |  |  |  |
| 7 | Giu 2020 | wt-Tom |  |  |  | 5 |  |  | 4 | 4 |
|  |  | R6/2-Tom |  |  |  | 5 |  |  | 4 | 7 |
|  |  | R6/2-hBP2 |  |  |  | 3 |  |  | 4 | 6 |

